# Simulated Microgravity Enhances Adipocyte Maturation and Glucose Uptake via Increased Cortical Actin Remodeling

**DOI:** 10.1101/2024.01.30.578049

**Authors:** Golnaz Anvari, Michael Struss, Evangelia Bellas

**Author notes:** ^$^ These authors contributed equally to this work (Golnaz Anvari, Michael Struss).

## Abstract

Adipose tissue (AT) regulates whole-body metabolism and is subject to various forces during movement, exercise, and during rest. Adipocytes are mechanically responsive cells, yet little is known about how the lack of mechanical loading may affect adipocytes and their function. To model the lack of mechanical loading, we exposed engineered AT constructs to simulated microgravity (sµg) conditions for 28 days. We found sµg enhanced lipid accumulation (lipogenesis) and lipid mobilization (lipolysis). Adipocyte maturation involves a phenotypic switch from actin stress fiber disruption and cortical actin formation. Sµg exposure increased cortical actin formation through mechanoresponsive signaling pathways involving Ras homolog family member A (RhoA) and Rho Associated Coiled-Coil Containing Protein Kinase 1 (ROCK1) downstream targets, cofilin and actin-related protein 2/3 (ARP2/3). Adipocytes cultured in sµg have increased glucose transporter type 4 (GLUT4) translocation to the cell membrane and insulin-stimulated glucose uptake, independent of the canonical Akt pathway. GLUT4 translocation to the cell membrane and insulin-stimulated glucose uptake was limited when we inhibited new formation of branched cortical actin using an ARP2/3 inhibitor, CK-666. This study demonstrated that sµg enhances adipocyte maturation via increased lipogenesis and lipolysis and cortical actin remodeling which further enhanced glucose uptake. Therefore, targeting these mechanosensitive pathways pharmacologically or simulating microgravity on earth as a non-pharmacological modality are novel approaches to improving adipocyte function and AT metabolism and possibly for treating related comorbidities such as type 2 diabetes and obesity.

## Introduction

Adipose tissue (AT) is a mechanically loaded tissue, and adipocytes, the primary cell type in AT, are exposed to various mechanical forces, such as tension, compression, and shear stresses/strains that occur in both static and dynamic activities such as sitting, lying down, or exercising ^1–3^. Previous in vitro and in vivo studies demonstrated that mechanical loadings could affect adipogenic differentiation through different mechanotransduction pathways confirming that adipocytes and their precursors are mechanoresponsive cells ^4,5^. Dynamic loading conditions, such as cyclic stretching, vibration, or static compression, inhibit adipogenic differentiation, while static stretching may enhance differentiation ^4–6^. Further, mechanical loading can be exerted on adipocytes in obese tissue due to AT fibrosis. For example, adipocytes cultured within an obese adipose tissue-derived decellularized matrix were morphologically different, producing a more flattened-out shape than adipocytes cultured in a lean tissue-derived decellularized matrix ^7^. The cell deformation observed in the obese decellularized matrix was comparable to cells experiencing 50% compression, a condition that decreased lipolysis and adipokine production and increased cytokines and expression of fibrosis mediators ^7^. However, no study has focused on how the lack of mechanical loading affects adipocytes and their function. In other cell types, such as fibroblasts and retinal epithelium cells, sµg causes actin cytoskeletal remodeling by inducing cortical actin formation ^8,9^. Cortical actin formation and remodeling are critical for adipogenesis and adipocyte maturation. This remodeling process involves actin stress fiber disruption and cortical actin formation to facilitate lipid accumulation ^10^.

Cortical actin formation includes actin branching regulated by the actin-related protein (ARP) complex. Further, cortical actin remodeling plays an essential role in regulating metabolic functions such as insulin-stimulated glucose uptake. Two pathways control glucose uptake in adipocytes, a mechanically independent, biochemical pathway regulated by insulin receptor substrate 1 (IRS-1) and protein kinase B (Akt) phosphorylation (pAkt) and a mechanical pathway that is dependent on cortical actin formation for shuttling glucose transporter type 4 (GLUT4) to the cell membrane ^11,12^. GLUT4 plays an important role in regulating overall glucose homeostasis ^13^. Therefore, inhibiting cortical actin remodeling and formation can prevent GLUT4 translocation and decrease glucose uptake without affecting downstream insulin signaling pathways such as IRS-1 and pAkt, confirming the critical role of dynamic cortical actin remodeling and formation in regulating glucose uptake ^14,15^.

To determine the effect of the lack of mechanical loading (i.e. gravity) on adipocyte cortical actin formation and remodeling and subsequent adipocyte function, we have exposed in vitro AT constructs, adipocytes encapsulated in spherical GelMA hydrogels, to simulated microgravity (sµg) for 28 days. In vitro three-dimensional (3D) AT models provide a reductionist model that enables us to systematically examine the interactions between adipocytes and their microenvironment ^16–24^. We found that adipocytes in sµg were morphologically different, including greater overall area and had greater lipogenesis (lipid accumulation) and lipolysis (free fatty acid release). Moreover, adipocytes in sµg had enhanced cortical actin remodeling and improved GLUT4 translocation to the cell membrane from increased ARP-mediated cortical actin formation, resulting in increased insulin-stimulated glucose uptake. Our findings were further validated by inhibiting cortical actin remodeling using the ARP protein complex inhibitor CK-666. Consistent with our results, inhibiting the ARP protein complex formation decreased insulin-stimulated glucose uptake due to less GLUT4 translocation to the cell membrane. Interestingly, when we separated the adipocytes into subpopulations by morphological features using flow cytometry, where smaller but greater number of lipid droplets represented a less mature adipocyte, and larger but fewer lipid droplets represented a more mature adipocyte, we found sµg exposure reduced the low maturity subpopulation and increased the high maturity subpopulation. ARP3 and insulin-stimulated glucose uptake also increased with adipocyte maturity.

Therefore, targeting mechanical pathways that regulate cortical actin remodeling using a pharmacological or novel non-pharmacological approach such as simulating microgravity on earth may be a new approach to improving glucose homeostasis, lipid metabolism and insulin sensitivity. Further studies would assess how changes in nutritional load, such as fatty acid supplementation, could alter adipocyte maturation and metabolism. Additionally, conditioned media from these studies could be used to assess how the adipocyte secretome may affect the function of other metabolic tissue types, such as muscle or bone, which are known to decrease in mass during spaceflight. Studies like this would aid in providing key insight into musculoskeletal changes observed in long-duration space flights ^25^.

## Results

### Adipocyte characterization in AT constructs

AT constructs were formed by encapsulating adipocytes within spherical gelatin methacryloyl (GelMA) hydrogels and cultured in 25 cm^2^ flasks (static condition, 1g) or HARV cartridges (simulated microgravity condition, sµg) for 28 days. More details about the experimental setup can be found in the Material and Methods section. Adipocyte morphology within AT constructs after a 28-day culture period was visualized via fluorescent confocal microscopy. Macroscopic images of whole AT constructs showed homogenously dispersed adipocytes within the AT constructs with robust fluorescent staining for BODIPY in both conditions (Fig. 2a). Fluorescent confocal images were taken at least 20 μm above the glass coverslip (Fig. 2b). Adipocyte size was significantly greater in sµg, indicating increased lipogenesis (Fig. 2c). Although adipocytes cultured in sµg have a greater overall size, the average size of individual lipid droplets and number of lipid droplets per adipocyte remained unchanged (Fig. 2c). Further, we assessed adipocyte related genes (Fig. 2d), including peroxisome proliferator-activated receptor gamma (*PPARG*), adiponectin (*ADIPOQ*), leptin (*LEP*), and lipogenesis/lipolysis related genes such as, perilipin-1 (*PLIN-1*), diglyceride acyltransferase (*DGAT*), and adipose triglyceride lipase (*ATGL*) (Fig. 2d, 2f, Supplementary Fig. S10, S11). *PLIN-1, DGAT, and ATGL* were significantly upregulated and *LEP* was significantly downregulated after 28 days in sµg compared to static controls (1g), while *PPARG* and *ADIPOQ* were not significantly affected (Fig. 2d). Lipid mobilization (lipolysis) is a critical adipocyte function; therefore, glycerol levels were quantified from banked media sampled weekly for the 28 days. Lipolysis increased after 14 days, with the highest rate of lipolysis on day 28 in sµg conditions (Fig. 2e). Together these results show that adipocytes cultured in sµg conditions had increased lipid metabolism, lipogenesis, and lipolysis compared to 1g conditions (Fig. 2g).

**Figure 1.**
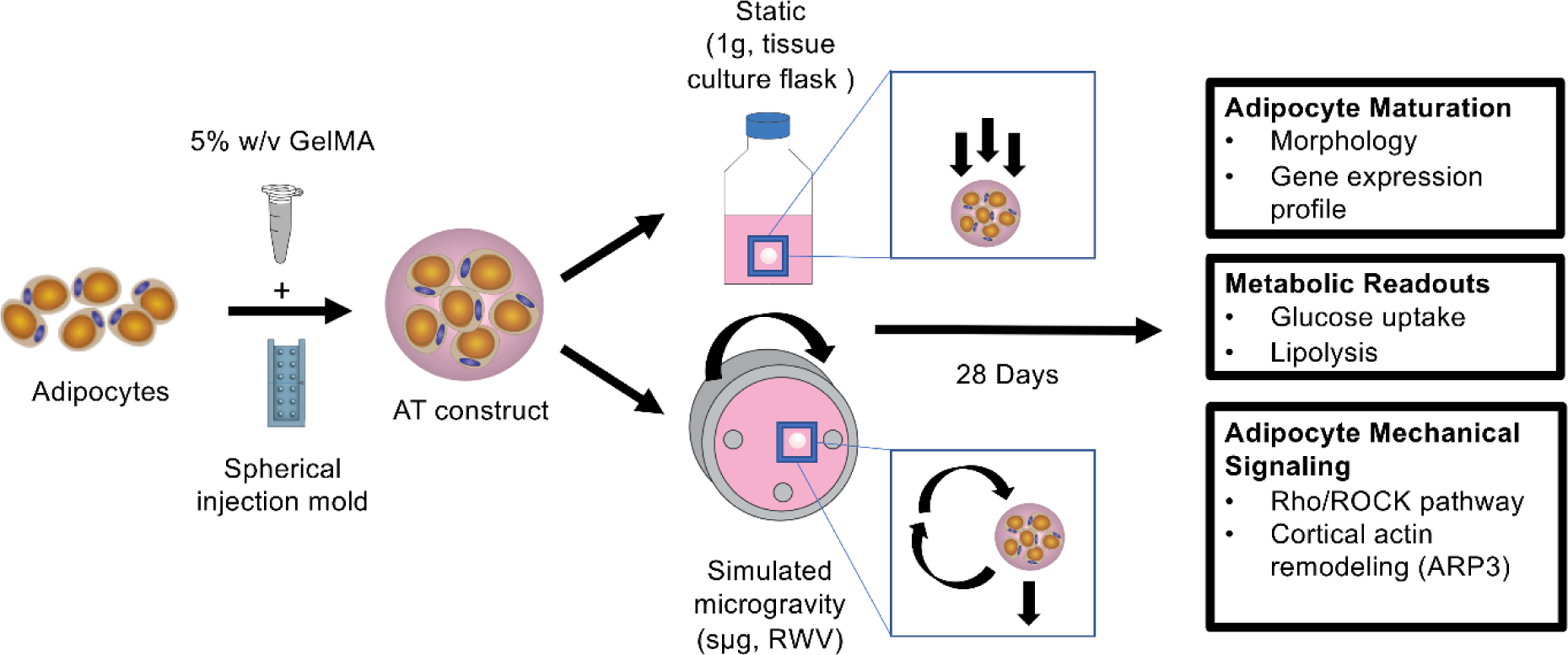
Experimental workflow. Adipocytes were encapsulated in 5% w/v GelMA using a spherical injection mold, these now will be referred to as AT constructs. AT constructs were kept in Static (1g) or Simulated microgravity (sµg) conditions for 28 days and sacrificed for endpoint analyses. Arrows illustrate the physical forces cells within the hydrogel experience as well as the moving direction of rotating wall vessel (RWV).

**Figure 2.**
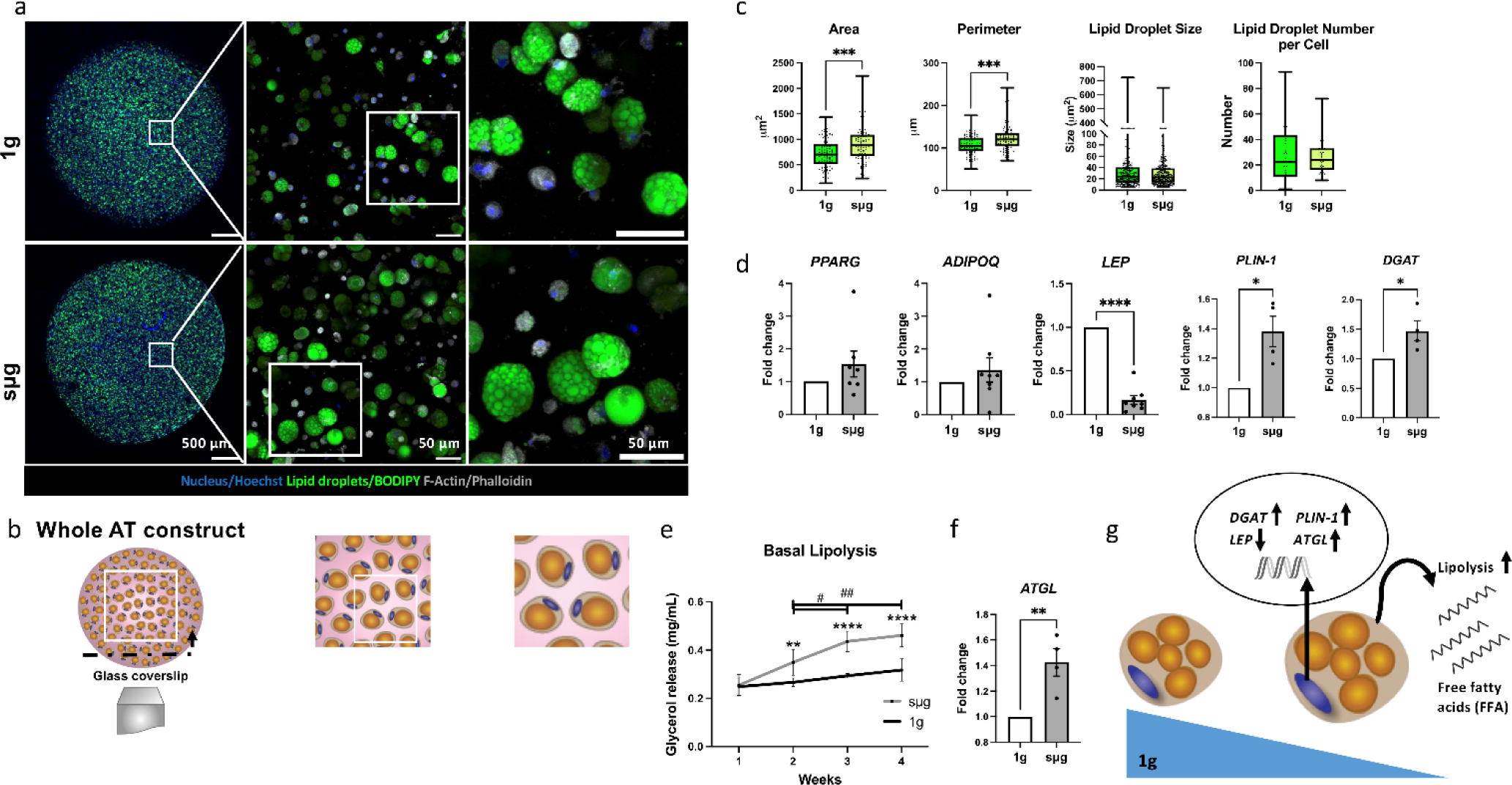
Adipocyte characterization in AT constructs. (a) Fluorescent confocal microscopy images (left panel, full AT construct, scale bar-500 µm, middle and right panel, scale bar-50 µm) of AT constructs after 28 days in 1g and sµg conditions. AT constructs were stained with Hoechst to visualize the nucleus, BODIPY to visualize the lipid droplets, and Phalloidin to visualize F-actin. (b) Cross-sectional fluorescent confocal microscopy images of AT constructs were taken at least 20 μm above the glass coverslip. (c) Adipocyte area and perimeter were significantly increased in sµg after 28 days (n=5 biological replicates, Box and Whiskers graphs are presenting min to max data with their median). Lipid droplet size (n=3 biological replicates, 9-11 cells quantified for each replicate) and lipid droplet number per adipocyte (n = 4 biological replicates, 6-10 cells quantified for each replicate) were not different across conditions after 28 days, Box and Whiskers graphs are presenting min to max data with their median). (d) Of the adipocyte genes, peroxisome proliferator-activated receptor gamma (*PPARG)*, adiponectin (*ADIPOQ),* and leptin (*LEP)*, *LEP* was significantly downregulated in sµg. Lipogenesis-related genes perilipin-1 (*PLIN-1)* and diacylglycerol acyltransferase (*DGAT*) were significantly upregulated in sµg (n=4-8 biological replicates). (e) Basal lipolysis levels were significantly increased in sµg (n=5 biological replicates). (f) Lipolysis-related gene, adipose triglyceride lipase (*ATGL*) was significantly upregulated in sµg (n=4 biological replicates). (g) Sµg increases lipid metabolism, lipogenesis, and lipolysis. Data are presented as means ± SEM. Comparisons between groups and statistical analysis were performed using two-way ANOVA with Tukey post hoc test, unpaired t-test or Mann–Whitney test with two-tailed p-values, *, ^#^ shows the statistical difference between different conditions (1g, sµg) at the same time point (*), and sµg at different time points (^#^) (*^, #^p < 0.05, **^, ##^p < 0.01,***p<0.001, ****p < 0.0001).

### Simulated microgravity increases cortical actin remodeling via increased ARP3 protein expression

Adipocytes are mechanoresponsive cells, and it known that sµg can induce cortical actin formation in other cell types ^8,9^. In adipocytes, cortical actin formation is a crucial for facilitating lipid accumulation. To investigate the effect of mechanical unloading (sµg) on adipocyte actin cytoskeletal remodeling, we assessed some key mechanotransduction pathways that regulate F-actin polymerization/depolymerization and cortical actin remodeling. Ras homolog family member A (*RHOA)* and Rho Associated Coiled-Coil Containing Protein Kinase 1 (*ROCK1)* gene expression were significantly upregulated in sµg conditions (Fig. 3a), but the total RhoA protein expression remained unchanged (Fig. 3b, Supplementary Fig. S1). Therefore, we evaluated one of the downstream targets of the RhoA/ROCK signaling pathway, cofilin and its phosphorylated inactive state, pcofilin, where it stabilizes actin rather than severs it. Exposure to sµg increased the pcofilin/cofilin ratio, confirming increased downstream RhoA/ROCK signaling pathway activity in sµg (Fig. 3c, Supplementary Fig. S2). To further investigate cortical actin remodeling, we checked actin branching complex molecules, ARP3 and ARP2 and found that ARP3 protein expression increased significantly in sµg, enhancing cortical actin remodeling (Fig. 3d, 3e, Supplementary Fig. S1,2). These results show that sµg enhances cortical actin remodeling via ARP3 protein expression (Fig. 3f).

**Figure 3.**
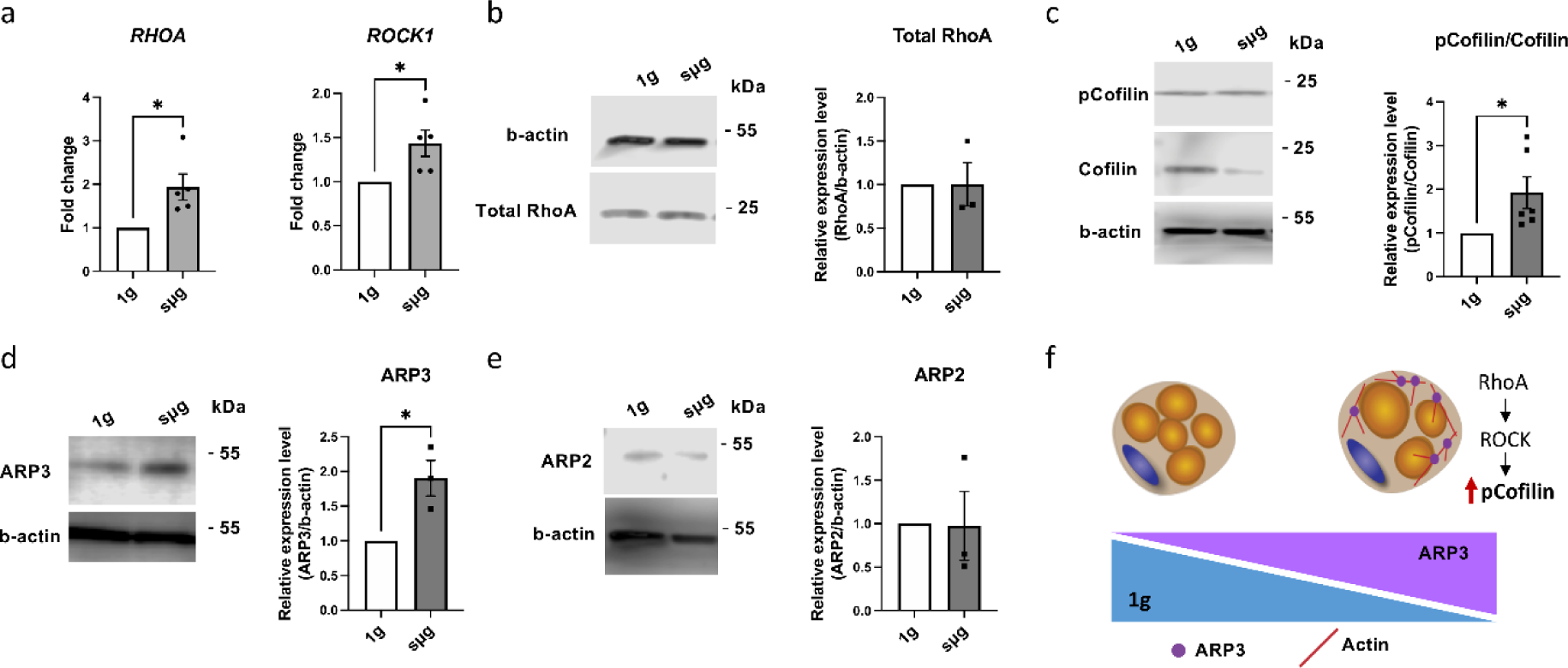
Simulated microgravity increases cortical actin remodeling via increased ARP3 protein expression. (a) Ras homolog gene family member A (*RHOA)* and Rho-associated coiled-coil kinases (*ROCK1)* gene expression was significantly increased in sµg (n=5 biological replicates). (b) Total RhoA protein expression was not affected in 1g and sµg conditions (n=3 biological replicates, the cropped blot is shown, the full-length blot is presented in Supplementary Fig. S1). (c) pCofilin/Cofilin protein expression was increased in sµg conditions (n=6 biological replicates, the cropped blot is shown, the full-length blot is presented in Supplementary Fig. S2). (d) Actin related protein 3 (ARP3) protein expression was significantly increased in sµg (n=3 biological replicates, the cropped blot is shown, the full-length blot is presented in Supplementary Fig. S1). (e) ARP2 protein expression was not affected in 1g and sµg conditions (n=3 biological replicates, the cropped blot is shown, the full-length blot is presented in Supplementary Fig. S2). (f) Simulated microgravity enhances adipocyte maturation via increasing ARP3 protein expression and enhancing cortical actin remodeling. Data are presented as means ± SEM. Comparisons between groups and statistical analysis were performed using an unpaired t-test with two-tailed p-values (*p < 0.05).

### Mature adipocyte subpopulation is significantly increased in simulated microgravity

Increased cortical actin formation enhances adipocyte maturation ^10^. Since we observed an increase in adipocyte size and overall lipid droplet area via fluorescent confocal microscopy (Fig. 2c), we further investigated these morphological differences using flow cytometry. To quantify these differences, we gated 3 subpopulations within the scatter plot based on increasing side scatter (SSC-A), Group I, II, III (Fig. 4a). Preadipocytes, identified by their low side scatter and weak BODIPY signal, were not included in our analysis (Supplementary Fig. S5, S6). Representative fluorescent confocal microscopy images for Groups I-III are shown in Fig. 4b. There was a significant increase in the number of adipocytes in Group III in sµg conditions compared to 1g (Fig. 4c). Together these results showed a significantly higher percentage of mature adipocytes, seen in Group III, in sµg after 28 days of culture.

**Figure 4.**
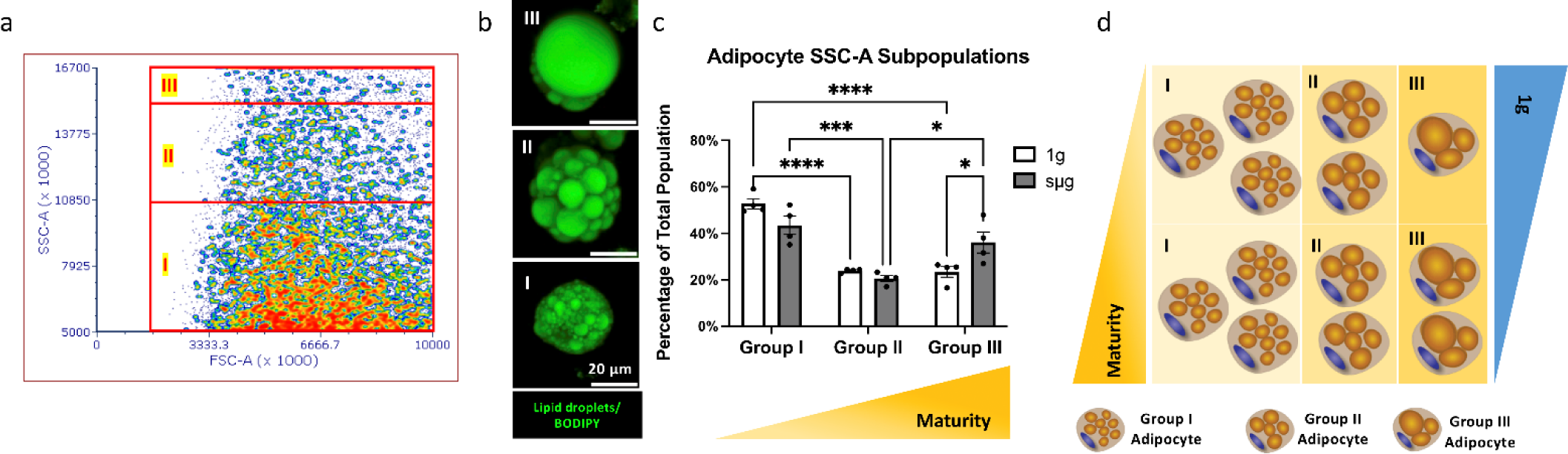
Mature adipocyte subpopulation is significantly increased in simulated microgravity. (a) Adipocytes were separated into three different subpopulations, referred to as Group I, Group II, Group III, based on increasing side scatter in flow cytometry scatter plots (n=4 biological replicates). (b) Representative fluorescent confocal microscopy images of Groups I-III (scale bar-20 µm). (c) The percentage of the mature adipocyte subpopulation, Group III, was significantly increased in sµg (n=4 biological replicates). (d) Adipocyte maturation increased in sµg conditions. Data are presented as means ± SEM. Comparisons between groups and statistical analysis were performed using two-way ANOVA with Tukey post hoc test (*p < 0.05, ***p < 0.001, ****p < 0.0001).

### Simulated microgravity enhances insulin-stimulated glucose uptake and GLUT4 translocation to the cell membrane

Cortical actin formation in adipocytes is not only necessary for adipocyte maturation but is also essential for glucose uptake via facilitating GLUT4 translocation to the cell membrane. To assess if increased cortical actin remodeling (Fig. 3c, 3d) and adipocyte maturation (Fig. 2c, 4c) affects glucose uptake in sµg, we quantified insulin-stimulated glucose uptake in AT constructs after 28 days in culture using a fluorescent analog of the glucose molecule, 2-NBDG. We found glucose uptake positively correlates with adipocyte maturation (Fig. 5a, 5b) and insulin-stimulated glucose uptake was significantly higher in sµg in the most mature adipocyte subpopulation, Group III (Fig. 5a). To confirm the relationship between this greater glucose uptake and GLUT4 translocation to the cell membrane, we quantified the ratio of GLUT4 fluorescent signal on the cell membrane to the cytoplasm. This ratio was significantly higher in sµg after insulin stimulation in Groups II and III (Fig. 5d). These results suggest that sµg enhances glucose uptake due to increased GLUT4 translocation to the cell membrane (Fig. 5e, Supplementary Fig. S12).

**Figure 5.**
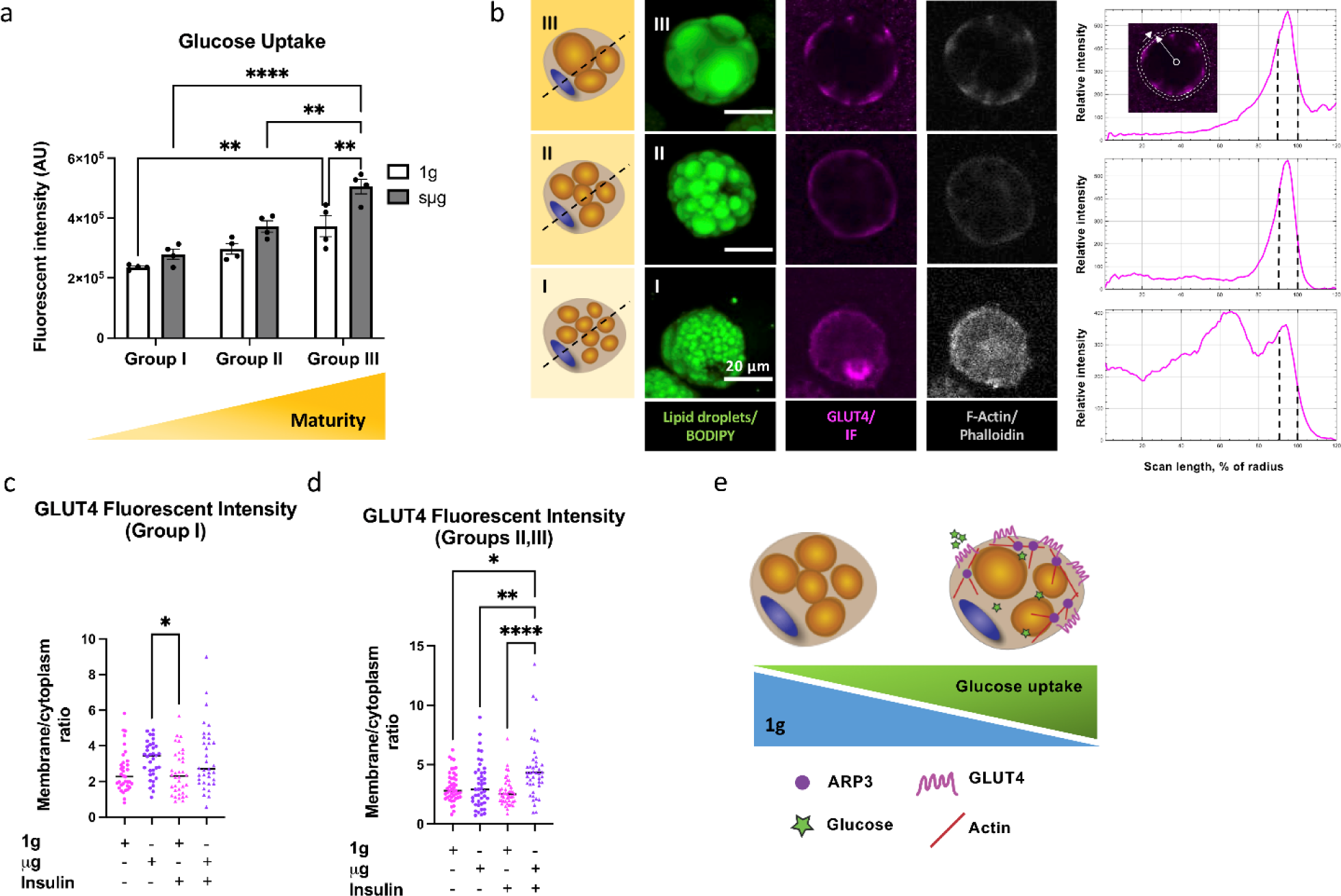
Simulated microgravity enhances insulin-stimulated glucose uptake and GLUT4 translocation to the cell membrane. (a) Insulin-stimulated glucose uptake was significantly greater in sµg conditions for all three groups (n=4 biological replicates). (b) Adipocyte maturation increases glucose transporter type 4 (GLUT4) translocation to the cell membrane and cortical actin formation; intensity plots are shown for GLUT4. To quantify GLUT4 fluorescent signal, adipocyte border was outlined using polygon line tool (inner white line) in ImageJ. Radial scan (white arrow) was set to 120% of the object radius (outer white line) from the object center (inner white circle). The average of fluorescent signal intensity between the black dashed lines in the intensity graph (90-100%) was quantified and divided to the average fluorescent intensity signal from (0-89%), images are illustrating the middle section of adipocytes. (c) GLUT4 membrane to cytoplasm ratio was not significantly different comparing both conditions for Group I (n=3 biological replicate, 10-12 cells were quantified for each replicate). (d) GLUT4 membrane to cytoplasm ratio was significantly increased in sµg for Groups II and III (n=4 biological replicate, 10-14 cells were quantified for each biological replicate). (e) Sµg enhances insulin-stimulated glucose uptake in adipocytes due to increased GLUT4 translocation to the cell membrane. Data are presented as means ± SEM. Comparisons between groups and statistical analysis were performed using two-way ANOVA or Kruskal-Wallis test with Tukey post hoc test or Dunn’s multiple comparisons test (*p < 0.05, **p < 0.01, ****p < 0.0001).

### Inhibiting cortical actin remodeling resulted in decreased glucose uptake in simulated microgravity

Next, we assessed the relationship between ARP3 protein expression and adipocyte maturity using flow cytometry. There was a positive correlation between ARP3 protein expression and adipocyte maturation (Fig. 6a). Earlier, we found that ARP3 protein expression is higher in sµg (Fig. 3d). Therefore, we hypothesized that increased ARP3 protein expression and cortical actin remodeling would increase insulin-stimulated glucose uptake in sµg. To check this hypothesis, we first investigated the Akt pathway in 1g and sµg after insulin stimulation. Sµg did not affect the ratio of pAkt/Akt protein expression (Fig. 6b). Then, to confirm the role of cortical actin remodeling in glucose uptake, we inhibited cortical actin formation by treating with an ARP3 inhibitor (CK-666). In the presence of CK-666 in sµg, insulin-stimulated glucose uptake was significantly decreased (Fig. 6c, Supplementary Fig. S7) by decreased GLUT4 translocation to the cell membrane (Fig. 6d, 6e) and cortical actin formation (Fig. 6f, 6g), confirming our hypothesis. While Ck-666 was shown to disrupt cortical actin remodeling, there was no difference in lipid droplet size after treatment with CK-666 (Supplementary Fig. S9). Together, these data support that ARP3 inhibition in sµg prevents cortical actin formation and results in decreased insulin-stimulated glucose uptake via decreased GLUT4 translocation to the cell membrane (Fig. 6h).

**Figure 6.**
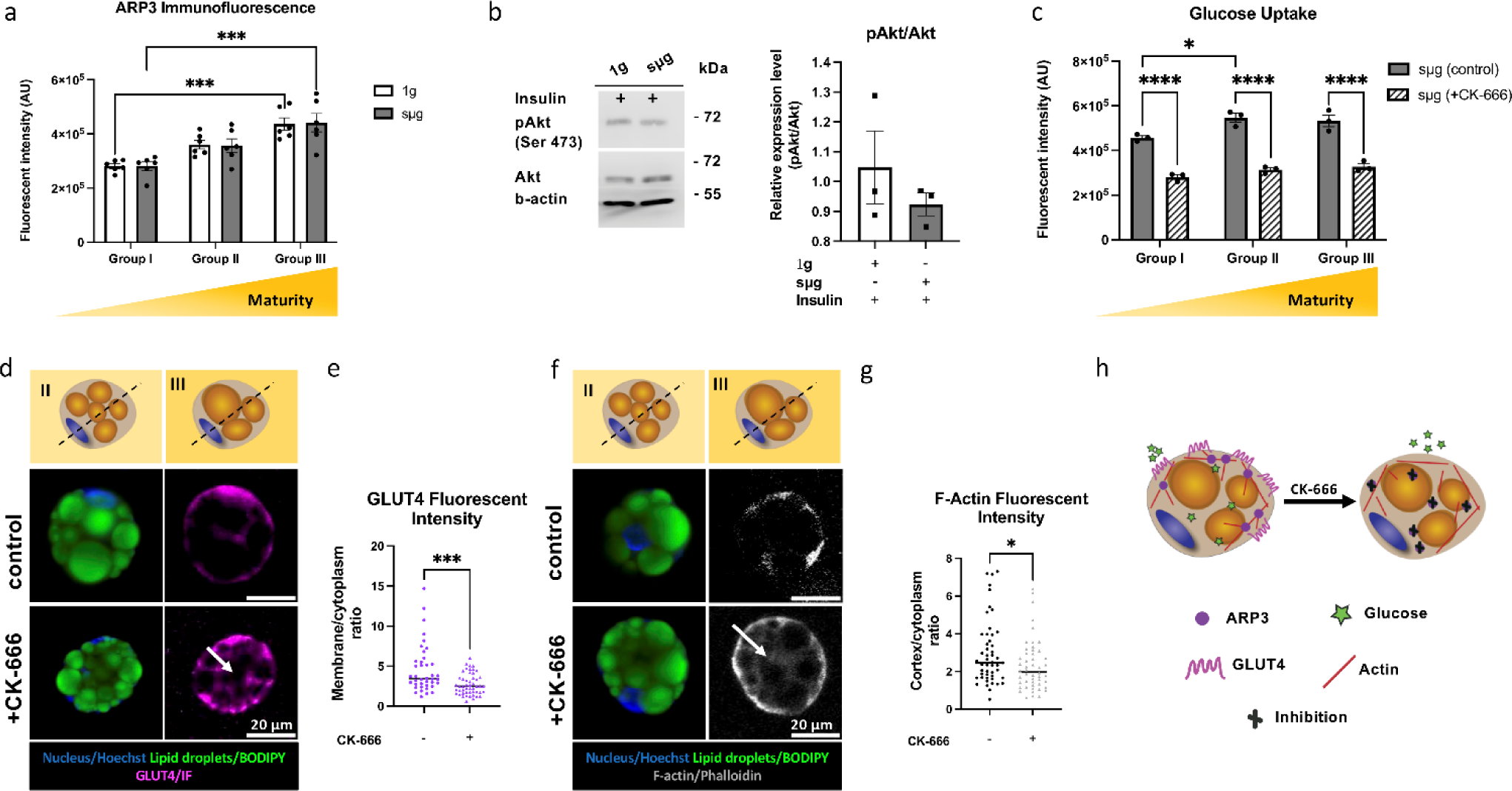
Inhibition of cortical actin remodeling decreased insulin-stimulated glucose uptake in simulated microgravity. (a) ARP3 expression was significantly increased in mature adipocytes (n=6 biological replicates). (b) pAkt/Akt protein expression was not affected in both conditions after insulin stimulation (n=3 biological replicates, data are normalized to 1g (-ins) condition, cropped blot is shown, the full-length blot is presented in Supplementary Fig. S3). (c) Insulin-stimulated glucose uptake was significantly downregulated in the presence of CK-666 in sµg conditions (n=3 biological replicates). (d) Representative cross-sectional images of adipocytes in the absence (control, top panel) or the presence of CK-666 (bottom panel) in sµg, white arrow points to the brighter cytoplasmic fluorescent signal for GLUT4 in the presence of CK-666 (scale bar-20 µm). (e) GLUT4 translocation to the cell membrane was significantly decreased in the presence of CK-666 (n=4 biological replicates, 10-12 cells were quantified for each replicate). (f) Representative cross-sectional images of adipocytes in the absence (top panel) or the presence of CK-666 (bottom panel) in sµg, white arrow points to the brighter cytoplasmic fluorescent signal for F-actin in the presence of CK-666 (scale bar-20 µm). (g) F-actin cortex to cytoplasm ratio was significantly decreased in the presence of CK-666 (n=3 biological replicates, 10-12 cells were quantified for each replicate). (h) ARP3 inhibition prevented cortical actin remodeling; therefore, GLUT4 translocation to the cell membrane and decreased insulin-stimulated glucose uptake. Data are presented as means ± SEM. Comparisons between groups and statistical analysis were performed using two-way ANOVA with Tukey post hoc test, Mann– Whitney test, or unpaired t-test with two-tailed p-values (*p<0.05, ***p < 0.001, ****p<0.0001).

## Discussion

We have summarized overall findings in Fig. 7. Adipocytes cultured in sµg for 28 days exhibited a more mature and metabolically active phenotype (Fig. 2, 4, 5). Moreover, there was a shift in subpopulations grouped by maturity, where there was a decrease in the low maturity group (Group I) and an increase in the high maturity group (Group III) (Fig. 4c). The master transcriptional regulator of adipogenesis, *PPARG*, and adipokine, *ADIPOQ* were not significantly upregulated (Fig. 2d) to match this phenotype, which may be attributed to both conditions cultured in the same adipogenic media containing a *PPARG* agonist, where biochemical stimulation overrides mechanical signaling ^26^. However, *LEP*, the gene encoding leptin, an adipokine which acts on the brain to signal satiety and is upregulated in obese patients, was downregulated (Fig. 2d). The decrease in serum leptin was also observed in a previous study of a mouse hindlimb unloading 28 day model of simulated microgravity ^27^. Leptin also plays multiple roles in adipocyte and whole-body metabolism, including in lipid metabolism. Exposure to increased free fatty acids (FFAs) has been shown to decrease *LEP* expression in rodent adipocytes ^28,29^. In this study, we found lipolysis to be increased (i.e. higher FFAs in media) with decreased LEP expression, similar to these studies showing FFA decreased with *LEP* expression. Additionally, we found *ATGL*, the gene encoding adipose triglyceride lipase, to be upregulated in sµg, supporting the finding of increased basal lipolysis (Fig. 2d). For lipogenesis-related genes, there was a significant upregulation in *PLIN-1* and *DGAT* (Fig. 2d). Perilipin is a lipid droplet coating protein that works as a barrier (when inactive, unphosphorylated) or a recruitment site (when active, phosphorylated) for lipases, while DGAT is an acyltransferase that catalyzes the formation of triglycerides ^30,31^. Therefore, adipocytes cultured in sµg could stimulate a more metabolically active, healthier adipocyte phenotype. Since brown adipocytes are often more metabolically active and the effect of sµg on mitochondrial function has been shown previously ^32^, we investigated the potential of induced browning in sµg. Browning related gene, uncoupling protein 1 (*UCP-1*) gene expression remained unchanged, while PPARG coactivator 1 alpha (*PGC-1α*) was significantly upregulated in sµg (Supplementary Fig. S10). *PGC-1α* is known to regulate lipid metabolism and fatty acid oxidation that further supports increased lipid metabolism in sµg ^33,34^.

**Figure 7.**
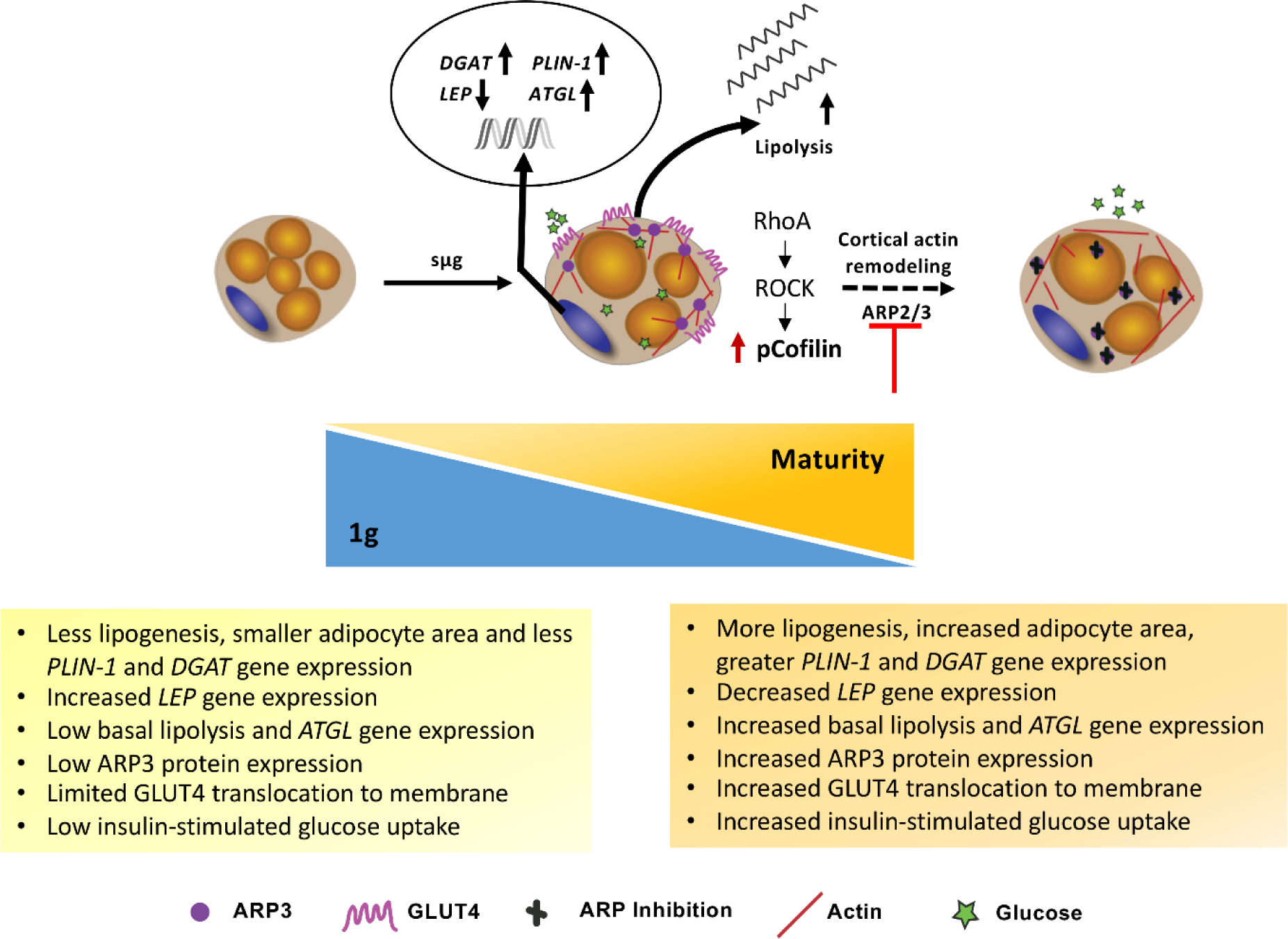
Summary overview. Simulated microgravity (sµg) enhances adipocyte lipid accumulation (lipogenesis) and lipid mobilization (lipolysis). Sµg increases cortical actin formation through mechanoresponsive signaling pathways involving downstream RhoA/ROCK effectors, Cofilin, and ARP2/3. Adipocytes in sµg have increased GLUT4 translocation to the cell membrane and insulin-stimulated glucose uptake, independent of canonical biochemical pathway including pAkt. Inhibition of branched cortical actin formation by ARP2/3 inhibitor, CK-666, reduced both GLUT4 translocation to the cell membrane and insulin-stimulated glucose uptake.

Since adipocyte maturation depends partly on cortical actin remodeling, we next investigated the mechanotransduction pathways that play an essential role in this remodeling. In part, actin remodeling in adipocytes is controlled through the RhoA/ROCK mechanical signaling pathway ^35^. RhoA/ROCK activity decreases during adipogenic differentiation to reduce actin filament formation and allow for lipid accumulation. However, RhoA/ROCK activity can also increase as adipocytes mature and become larger ^6,36^. Cofilin is one of the downstream targets in the RhoA/ROCK signaling pathway and is responsible for F-actin depolymerization. However, when cofilin is phosphorylated through the active RhoA/ROCK signaling pathway, it leads to F-actin stabilization. A previous study indicated that the F/G actin ratio has an important role in cortical actin formation, meaning that the higher F/G actin ratio can be translated to accelerated F-actin transition and increased cortical actin formation ^37^. This is partly controlled through actin severing molecules including cofilin; therefore, we concluded that increased pcofilin/cofilin ratio in sµg can be defined as increased F/G actin ratio and increased cortical actin formation (Fig. 3c, Supplementary Fig. S2) ^37^. Further, cortical actin remodeling depends on two processes: actin filament formation and actin branching. Branched actin filament formation, cortical actin, is regulated by ARP2/3 complex ^38^. Previous studies demonstrated that ARP3 localization and protein expression were affected in the early stage of obesity as well as during adipocyte maturation. In the early stage of differentiation, ARP3 is mainly accumulated around the cell cortex to facilitate cortical actin formation ^10^. Then, during adipocyte maturation, increased cortical actin remodeling is needed to support lipid accumulation resulting in more ARP2/3 protein expression^36^. Therefore, we concluded that in sµg increased ARP3 protein expression favors cortical actin stabilization and remodeling (Fig. 3d).

Cortical actin remodeling also plays a dynamic role in insulin-stimulated glucose uptake. In adipocytes, insulin-stimulated glucose uptake is dependent on GLUT4 translocation to the cell membrane, and cortical actin assembly is involved in GLUT4 storage vesicle docking and tethering to the cell membrane ^39^. Previous studies have shown that disrupting actin filament formation can reduce insulin-stimulated glucose uptake without affecting the Akt signaling pathway, thus confirming the essential role of these cortical actin filaments in shuttling GLUT4 vesicles to the cell membrane ^13^. There was no change in insulin-stimulated glucose uptake when switching from sµg to 1g after insulin starvation (Supplementary Fig. S8), however there was a decrease in GLUT4 translocation to the membrane after inhibiting the ARP3 protein complex, in agreement with this previous literature ^23^.

Overall, we found that in sµg conditions, adipocyte maturity was enhanced as visualized by significantly larger lipid droplet area. Enhanced maturity was further confirmed by a shift in subpopulation distribution towards a more mature adipocyte phenotype. Adipocytes have an increased rate of lipogenesis (lipid accumulation) and lipolysis (free fatty acid release) in sµg conditions. Additionally, GLUT4 translocation to the membrane and insulin-stimulated glucose uptake was enhanced in sµg via cortical actin remodeling (Fig. 7).

Adipocyte mechanical properties including actin cytoskeletal remodeling can be affected under different conditions such as obesity leading to a dysfunctional AT. Previously, our group has confirmed adipocytes to be mechanoresponsive cells by investigating actin remodeling in the presence of different obesity related microenvironmental cues such as hypoxia or altered collagen stiffness and architecture ^40,41^. Our findings showed that adipocytes within the obese microenvironment show a high incidence of persistent actin stress fibers, hindering adipocyte maturation or preadipocyte differentiation ^40,41^. Obesity can result in other comorbidities such as type 2 diabetes, a dysregulation of insulin sensitivity and glucose homeostasis. Therefore, targeting cortical actin remodeling pharmacologically or using exposure to sµg as a non-pharmacological therapy could improve glucose homeostasis and insulin sensitivity in type 2 diabetic and obese patients by enhancing adipocyte metabolic function and maturation via cortical actin remodeling. AT supports other tissues through endocrine crosstalk, and poor management of AT metabolism, in cases such as obesity, has been linked to higher incidences of arthritis in weight-bearing and non-weight-bearing joints ^42,43^. Thus, sµg may not be limited to enhancing AT metabolism but has implications in managing underlying diseases that result from AT dysregulation.

## Materials and Methods

### High aspect ratio rotating vessel (HARV) cartridge fabrication and sterilization

HARV cartridges were assembled and sterilized as described ^44^. In brief, cylindrical HARV cartridges were comprised of two external 3-inch outer diameter plates of polymethyl methacrylate (PMMA) and a hollow central PMMA plate with a 2-inch inner diameter which formed the 10 mL cylindrical culture chamber. A fused Nylon/Tegaderm™ membrane was placed between the central and bottom PMMA plates to allow for gas exchange. Silicon gaskets were placed between layers to ensure a tight seal. Once all layers were aligned, they were secured with stainless steel bolts to create a watertight and gas-permeable culture chamber. Luer-lock injection ports were added to the front of the cartridge for media exchange and AT construct loading. Prior to culture, HARV cartridges were sterilized with sodium hydroxide and 1X phosphate buffer saline (PBS, Fisher, Cat # BP3994) containing 10% fetal bovine serum (FBS, Sigma-Aldrich, Cat # 12306C) and 2% penicillin/streptomycin (P/S, HyClone, Cat # SV30010).

### Gelatin methacryloyl (GelMA) preparation

10 grams of gelatin (Sigma-Aldrich, type A, 300 bloom, Cat # G2500) were dissolved in 100 mL of 0.25 M sodium bicarbonate buffer (Fisher, Cat # 144-55-8) at 50°C on a stirring hot plate until completely dissolved. Once dissolved, 2.5 mL (2.5% v/v) Methacrylic anhydride (Sigma-Aldrich, 94%, Cat # 760-93-0) was added dropwise at a flow rate of 0.6 mL/minute to the gelatin solution and allowed to react for 1 hour. During the reaction, the pH was adjusted to 9 and protected from light. The gelatin methacryloyl (GelMA) solution was diluted at 1:4 in prewarmed 1X PBS and placed in dialysis tubes (Sigma-Aldrich, 12-14 kDa MWCO, Cat # 21-152-14). The GelMA solution was dialyzed for 3 days against 3L of 1X PBS followed by 3 days against 3L of deionized water. All dialysis was performed at 55°C on a stirring hot plate and protected from light. After dialysis, the GelMA solution was sterile and filtered through a 0.22 µm filter, aliquoted into sterile tubes, snap-frozen, and lyophilized. The lyophilized GelMA was stored at −80°C until use.

### Spherical injection mold fabrication and sterilization

The spherical injection mold was designed using CAD software (OnShape) and 3D printed using the Objet Connex2 350 3D printer (Stratasys) with Vero glossy finish photo resin. In brief, Polydimethylsiloxane (PDMS; Sylgard 184, Dow Corning, Cat # 2065622) was mixed at a 10:1 ratio of base polymer to curing reagent. The mixture was cast in the 3D printed template at 100°C for 45 minutes to crosslink the PDMS. The spherical injection molds were removed from the 3D printed template and soaked in 100% isopropanol for 4 hours to remove any contaminants from the 3D printed resin or any un-crosslinked PDMS oligomers. The spherical injection molds were soaked overnight in double-distilled water (ddH_2_O) to remove excess isopropanol. The spherical injection molds were autoclaved prior to use.

### Two-dimensional (2D) cell culture and adipogenic induction

Human bone marrow-derived mesenchymal stem cells (hMSCs) were isolated from healthy adult male bone marrow, purchased from Lonza Walkersville Inc. (Walkersville, MD, Cat# 1M-105). HMSCs were expanded in low glucose Dulbecco’s Modified Eagle’s medium (DMEM, Corning, Cat # 10014CM) supplemented with 10% FBS and 1% P/S in 2D until confluency. Adipogenic induction media was added when hMSCs reached confluency at passage 4. The adipogenic induction media was composed of Dulbecco’s Modified Eagle’s medium/ Ham’s F-12 (DMEM/F12, Corning, Cat # 10092CM) supplemented with 3% FBS, 1% P/S, 2 µM rosiglitazone (Cayman Chemical, Cat # 71740100), 500 µM 3-isobutyl-1-methylxanthine (Thermo Scientific, Cat # 28822-58-4), 1 µM dexamethasone (Thermo Scientific, Cat # 50-02-2), 33 µM biotin (Thermo Scientific, Cat # 58-85-5), 17 µM D-calcium pantothenate (TCI America, Cat # 137-08-6), and 20 nM of insulin (Sigma-Aldrich, Cat # I9278). The hMSCs were differentiated for 7 days prior to encapsulation. Cells were maintained at 37°C and 5% CO_2_.

### Three-dimensional (3D) cell encapsulation in spherical injection mold

After 7 days of adipogenesis, adipocytes were detached via trypsin (Gibco) and resuspended at 8 million/mL in 5% (w/v) GelMA in 1X PBS with 0.05% (w/v) Lithium phenyl-2,4,6-trimethyl-benzoyl phosphinate (LAP, Allevi). This adipocyte containing hydrogel precursor solution was then pipetted into the spherical injection mold (30 µL per sphere in the spherical injection mold) and photocrosslinked with a 405 nm light (15 mW) for 2 minutes.

### Static (1g) and simulated microgravity (sµg) culture

The adipocyte containing GelMA spherical hydrogels (referred to now as *AT constructs*) were transferred to 25 cm^2^ flasks (static condition, 1g) or HARV cartridges (simulated microgravity condition, sµg). Each flask and cartridge contained 6 AT constructs in 10 mL of adipogenic media. The HARV cartridges were loaded onto a custom-built rotating wall vessel (RWV) bioreactors modeled after the Synthecon RCCS-4DQ or the Synthecon RCCS bioreactor itself. The commercial and custom-built bioreactors yielded no significant differences in outcomes. The bioreactor was set to a rotation speed of 25 rpm to keep the AT constructs in a simulated free-fall like state. Adipogenic media was replenished every other day, and AT constructs were maintained in either static (1g) or simulated microgravity (sµg) conditions for 28 days (Fig. 1).

### Lipolysis

Media was banked every 7 days for 28 days. Basal lipolysis levels were assessed by measuring the glycerol concentration in the banked media as determined by the Sekure Triglyceride Reagent Kit (Sekisui Diagnostics, Cat # 23660) and measured using the Infinite M200 Pro plate reader (TECAN). All conditions were performed in technical duplicates.

### Gene expression

Two pooled AT constructs per condition were stored in Trizol (Invitrogen, Cat # 15596026) at −80°C. RNA was isolated using the chloroform (Acros Organics, Cat # 67-66-3) phase separation method and precipitated in 50% isopropanol (Fisher Chemical, Cat # 67-63-0). The precipitated RNA was washed twice with 75% ethanol and resuspended in diethylpyrocarbonate (DEPC)-treated water (Invitrogen, Cat # AM9920). Total RNA reverse transcription and Real-time PCR (RT-PCR) was performed with High-Capacity cDNA Reverse Transcription Kit (Applied Biosystems, Cat # 4368814) and Power Up SYBR Green Master Mix (Applied Biosystems, Cat # A25776), respectively. Primers were custom designed (Integrated DNA Technologies). Samples were run in technical triplicates using an Eppendorf Mastercycler and StepOnePlus PCR system (Applied Biosystems). Fold change in relative gene expression levels was determined using the 2^-ΔΔCt^ method.

### Protein expression

Protein extracts from 2 pooled AT constructs were obtained by lysing the adipocytes in radioimmunoprecipitation assay (RIPA) buffer (supplemented with a phosphatase inhibitor cocktail containing 250 mM sodium fluoride, 50 mM sodium orthovanadate, 50 mM Sodium pyrophosphate decahydrate, and 50 mM β-glycerophosphate (Thermo Scientific, Cat # 78440). Proteins were separated by gel electrophoresis using 8% or 12% SDS-PAGE gel and transferred onto a nitrocellulose membrane (Immobilon-P, Cat # IPVH00005). After blocking and probing with antibodies, detection was performed using horseradish peroxidase a conjugated anti-rabbit secondary antibody (1:3000, Cat# 170-6515, RRID: AB_11125142) or an anti-mouse secondary antibody (1:3000, Cat# 170-6516, RRID: AB_11125547) and enhanced chemiluminescence reagent (VWR, Cat # 490005-018). The signal was visualized using a C-DiGit Blot Scanner (Li-Cor). For protein detection, the following primary antibodies (Cell Signaling Technology) were used: rabbit anti-actin related protein 3 (ARP3) (1:1000, Cat# 4738, RRID:AB_2221973), rabbit anti-ARP2 (1.500, Cat# 3128, RRID:AB_2181763), rabbit anti-ras homolog family member A (RhoA) (1:1000, Cat# 2117, RRID:AB_10693922), mouse anti-Akt (1:1000, Cell Signaling, Cat# 2920, RRID:AB_1147620), phosphorylated Akt (pAkt-Ser473) (1:1000 Cat# 4060, RRID:AB_2315049), rabbit anti-cofilin (1:500, Cat# 3318, RRID:AB_2080595), rabbit anti-phosphorylated cofilin (1:500, pcofilin-Ser3, Cat# 3311, RRID:AB_330238), and rabbit anti-beta actin (1:1000, Cat# 4970, RRID:AB_2223172), as a housekeeping protein.

### Visualization by immunofluorescence and cell staining

For immunofluorescence analysis, AT constructs were fixed with 4% (v/v) paraformaldehyde (Thermo Scientific, Cat # 28908) for 1-3 hours at room temperature (RT) and then permeabilized with 0.1% (v/v) Triton X-100 (Sigma-Aldrich, Cat # 9036-19-5) for 1 hour before blocking with 3% (w/v) bovine serum albumin (BSA, Fisher Scientific, Cat # BP9703100). To detect ARP3 and GLUT4, the following primary antibodies were used: rabbit anti ARP3 (1:100, Cell Signaling Technology, Cat# 4738, RRID: AB_2221973), and mouse anti-GLUT4 (1:100, Cell Signaling Technology, Cat# 2213, RRID: AB_823508). AT Constructs were co-stained with Hoechst 33342 (1:1000, Thermo Scientific, Cat # 62249), BODIPY 493/503 (1:100, Invitrogen, Cat # D3922), and Alexa Fluor 647 phalloidin (1:200, Invitrogen, Cat # A22287) for 45 minutes at RT to visualize nuclei, lipid droplets, and actin, respectively, and stored in 1X PBS at RT until imaged.

### Fluorescent confocal microscopy and image analysis

Images were acquired with a laser scanning confocal mode of the hybrid confocal multiphoton microscope (Olympus FluoView FV1200) using a 10X (UPLXAPO10X, 0.4 NA, Olympus) or 30X (UPLSAPO30XSIR, 1.05 NA, Olympus) oil immersion inverted objectives with four laser units (405, 488, 543, and 635 nm.), and four photomultipliers (PMT) detectors. Z-stacks (thickness ∼ 50 µm, step-size: 5 µm, 12.5 µs/pixel, 1024×1024) were captured at least 20 µm above the glass coverslip to ensure visualized cells were in a 3D environment and not in contact with the glass coverslip. When appropriate, brightness and contrast were applied equally to all conditions to improve signal and reduce background. Adipocyte size (area and perimeter), lipid droplet size, and the number of lipid droplets per adipocyte were quantified by separating the BODIPY fluorescent channel from confocal image stacks and slices from the stacks were z-projected using the Max Intensity method. To quantify adipocyte size, individual adipocyte images were converted to binary masks, and holes were filled manually. Area and perimeter were calculated using the “ShapeFilter” measurement tool ^45^. To quantify lipid droplet size, the Lipid Droplets Tool (http://dev.mri.cnrs.fr/projects/imagej-macros/wiki/Lipid_Droplets_Tool) was used. Lipid droplets without discernable edges were omitted from the analysis and only individual lipid droplet size measurements were analyzed. The number of lipid droplets per adipocyte were counted manually by counting only the lipid droplets with discernable edges. All measured values were converted to µm^2^. For quantifying GLUT4, cells were outlined using polygon selection, and an intensity profile was made for each cell using a multi-clock scan macro in ImageJ ^46^. The cell membrane to cytoplasm ratio for GLUT4 was calculated by averaging the intensity value for the cell membrane (90-100%) divided by the average intensity value for the cytoplasm (0-89%). The same strategy was used to quantify the ratio of F-actin in the cortex to the cytoplasm.

### Insulin-stimulated glucose uptake and ARP3 inhibition

To perform insulin-stimulated glucose uptake assays, AT constructs were first insulin starved for 6 hours. After 6 hours, media was replaced with media containing insulin and 20 µM fluorescent analog of glucose, 2-(*N*-(7-Nitrobenz-2-oxa-1,3-diazol-4-yl)Amino)-2-Deoxyglucose (2-NBDG, Invitrogen, Cat # N13195) and incubated for 30 minutes in their respective culture conditions. After 30 minutes, AT constructs were washed 3 times with 1X PBS and fixed with 4% (v/v) formaldehyde for 1 hour at RT, followed by 3 washes in 1X PBS. For ARP3 inhibition assays, samples were exposed to 100 µM CK-666 (Tocris Bioscience, 39-501-0) in dimethyl sulfoxide (DMSO, Sigma Aldrich. Cat # 67-68-5) during insulin starvation and stimulation.

### Flow cytometry sample preparation and analysis

AT constructs were stained with 2-NBDG, BODIPY, or ARP3 immunostaining as described above for their respective quantification. All AT constructs were stained with a DRAQ5 fluorescent probe (1:1000, Thermo Scientific, Cat # 62254) for 1 hour and washed 3 times in 1X PBS prior to enzymatic digestion. The AT constructs were digested in 100 µL of 35 mg/mL collagenase type I (Gibco, Cat # 17100017) for 30 minutes at 37°C. AT constructs in the collagenase solution were further dissociated by pipetting up and down and resuspended in 1X PBS to a final volume of 300 µL. Flow cytometry was performed using an Accuri C6 flow cytometer (BD Biosciences), and data were acquired and analyzed using BD CSampler analysis software (BD Biosciences). 300 µL of cell suspension was collected per sample, and events in the DRAQ5-FL4-A fluorescent channel (647 nm excitation) below 10^3 were excluded from data collection to minimize events from debris. Post collection, events from the debris were further removed by adjusting the FL4-A gate, deleting events below 2,000,000 forward scatter area (FSC-A), and removing events with a width greater than 250 or less than 50 (AU) (Supplementary Fig. S4). Three distinct subpopulations were gated based on SSC-A (Group I: 2 million – 10 million FSC-A, 5 million – 10 million SSC-A, Group II: 2 million – 10 million FSC-A, 10 million – 15 million SSC-A, Group III: 2 million – 10 million FSC-A, 15 million – 16.7 million SSC-A), and mean fluorescence intensity was recorded from the FL1 channel (488 nm) ^47–50^. Scatter plots and histograms were created using FCS Express 7 flow cytometry software (De Novo Software).

### Statistics

GraphPad Prism 8 software was used to perform all statistical analyses. All experiments were conducted with at least three biological replicates and 10-20 cells per biological replicate for image quantification. The distribution of each data set was analyzed, and the D’Agostino-Pearson test (α=0.05) was performed to test for normality. Statistical comparisons between two experimental groups were performed using two-tailed Student’s t-tests or Mann-Whitney test for non-normally distributed data when appropriate. Comparisons among multiple groups were performed using two-way analysis of variance (ANOVA) or Kruskal-Wallis test with Tukey post hoc testing. All graphs are presented as mean ± standard error of the mean (SEM) unless otherwise stated. Significance was determined according to *p *<* 0.05, **p ≤ 0.01, ***p ≤ 0.001, and ****p ≤ 0.0001.

## Acknowledgments

The authors would like to acknowledge funding support from NASA Space Biology grant 80NSSC19K0427 to E.B. The authors would like to thank Michael Phelan and Joseph Licata for their assistance in bioreactor cartridge assembly and setup.

## Author contributions

G.A. and M.S. performed the experiments and data analysis. G.A., M.S., and E.B. designed the experiments, interpreted the results, and wrote the paper. All three authors reviewed the manuscript.

## Competing interests

The authors declare no competing interests.

## Data availability

The datasets generated and analyzed during the current study are available from the corresponding author upon reasonable request.

## Notes

### Competing Interest Statement

The authors have declared no competing interest.

