## Supplementary information for "Simulated Microgravity Enhances Adipocyte Maturation and Glucose Uptake via Increased Cortical Actin Remodeling"

* Corresponding Author

**This file includes:**

Supplementary Figs. S1 to S12

Supplementary Tables. S1 to S37

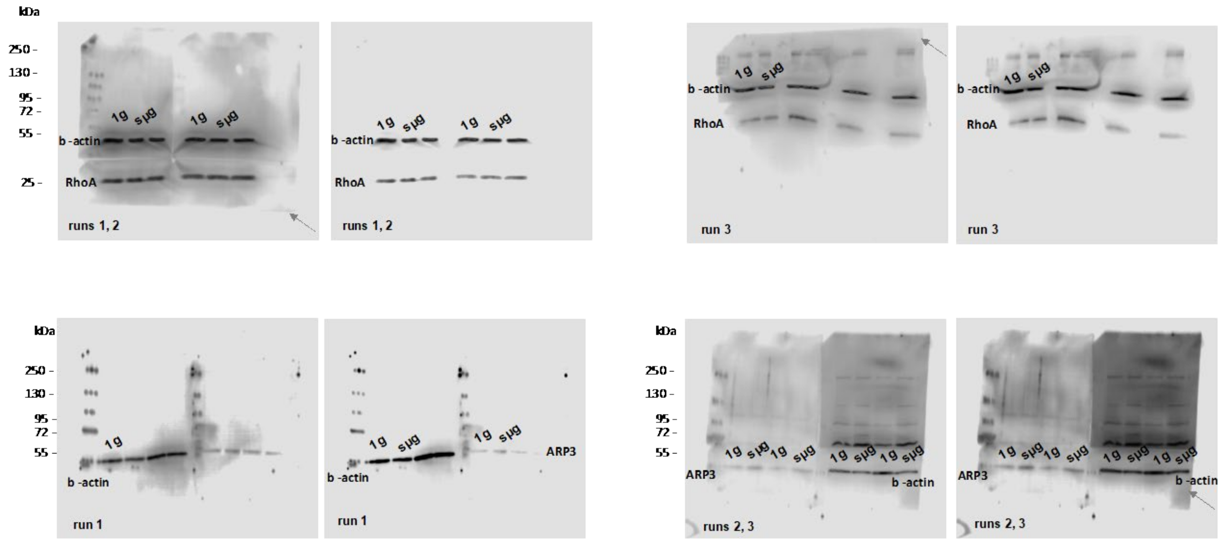

**Supplementary Figure S1. Full length blots for RhoA and ARP3 protein expression.** RhoA and ARP3 protein expression in 1g and sμg conditions, with two different intensities for each biological replicate (n=3 biological replicates). Brightness is not adjusted for the blots and different image displays are being presented to visualize the edge of the blotting membrane (grey arrows).

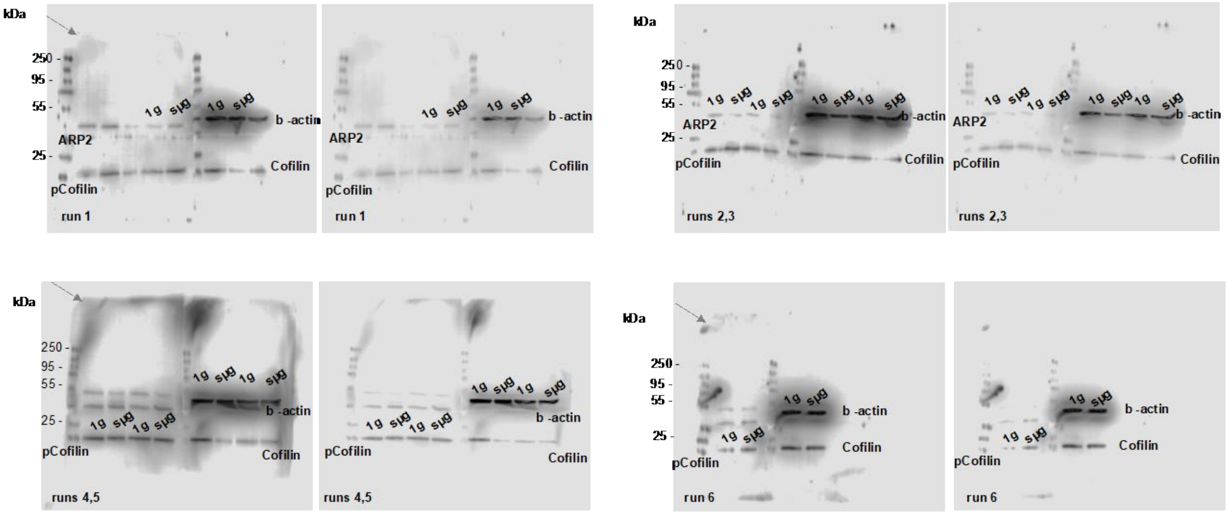

**Supplementary Figure S2. Full length blots for ARP2, pCofilin, and Cofilin protein expression.** ARP2, pCofilin and Cofilin protein expression in 1g and sμg conditions, with two different intensities for each biological replicate (n=3 biological replicates for ARP2 and n=6 biological replicates for Cofilin and pCofilin). Brightness is not adjusted for the blots and different image displays are being presented to visualize the edge of the blotting membrane (grey arrows).

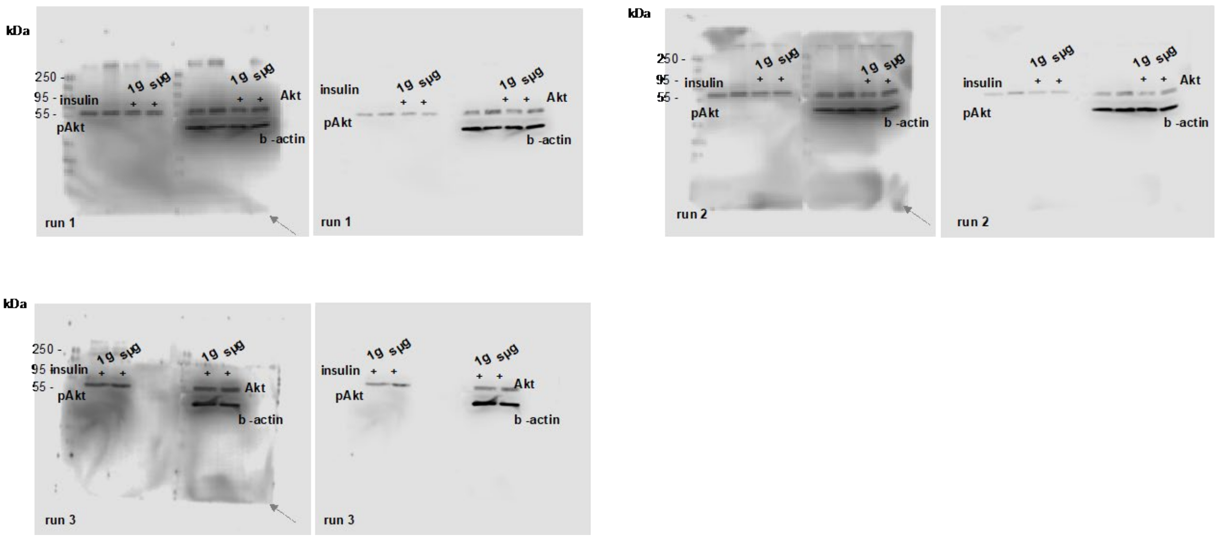

**Supplementary Figure S3. Full length blots for Akt, and pAkt.** Akt and pAkt protein expression in 1g and sμg conditions, with two different intensities for each biological replicate (n=3 biological replicates, top panel). Brightness is not adjusted for the blots and different image displays are being presented to visualize the edge of the blotting membrane (grey arrows).

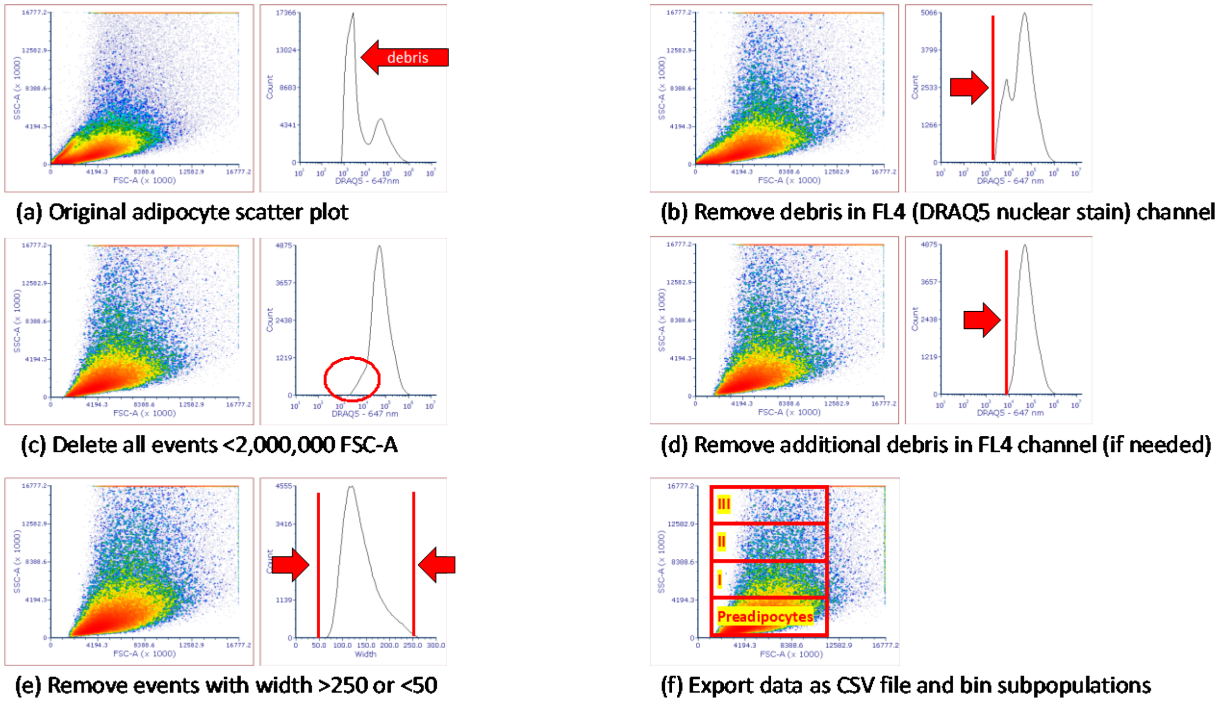

**Supplementary Figure S4. Flow Cytometry Gating Strategy for Adipocytes.** (a) Any events below 10^3 in the FL4-A (647 nm, DRAQ5, nuclear stain) channel are not collected to minimize events from debris. (b) The gate on the FL4-A channel is adjusted to the right. (c) Any events with <2,000,000 FSC-A are deleted further remove debris. (d) The FL4-A gate may need further adjustment if a conservative gate was used initially in (a). One distinct peak should remain, and this is our adipocyte population. (e) Events with a width >250 or < 50 are removed to avoid adipocyte clumps or doublets. (f) The data is exported as a CSV and events are binned into subpopulations (Group I: 2 million – 10 million FSC-A, 5 million – 10 million SSC-A, Group II: 2 million – 10 million FSC-A, 10 million – 15 million SSC-A, Group III: 2 million – 10 million FSC-A, 15 million – 16.7 million SSC-A).

a

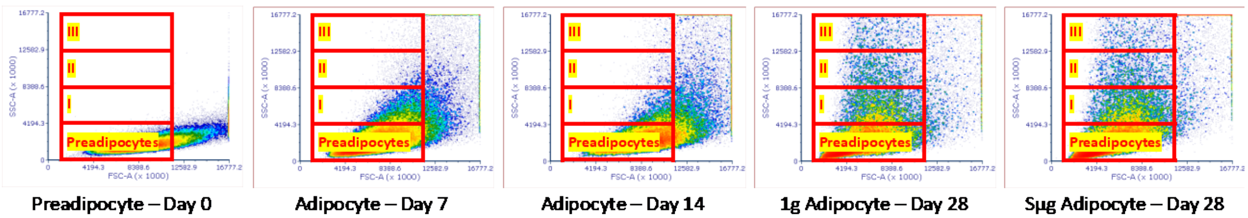

b

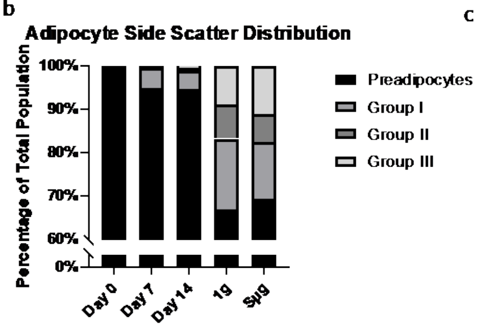

c

**Percentage of Population**

| | Preadipocytes | Group I | Group II | Group III |
|---|---|---|---|---|
| Day 0 | 99.99% | 0.00% | 0.00% | 0.00% |
| Day 7 | 94.91% | 4.59% | 0.42% | 0.08% |
| Day 14 | 94.69% | 4.25% | 0.65% | 0.42% |
| 1g (Day 28) | 70.59% | 15.34% | 7.03% | 7.03% |
| Sμg (Day 28) | 72.22% | 11.84% | 5.72% | 10.22% |

**Supplementary Figure S5. Using Side Scatter to Bin Adipocyte Subpopulations.** (a) Scatter plots of adipocytes at different time points of differentiation (day 0, day 7, day 14, day 28 static, day 28 sμg). (b,c) Percentage of each adipocyte subpopulation in the total adipocyte population.

a

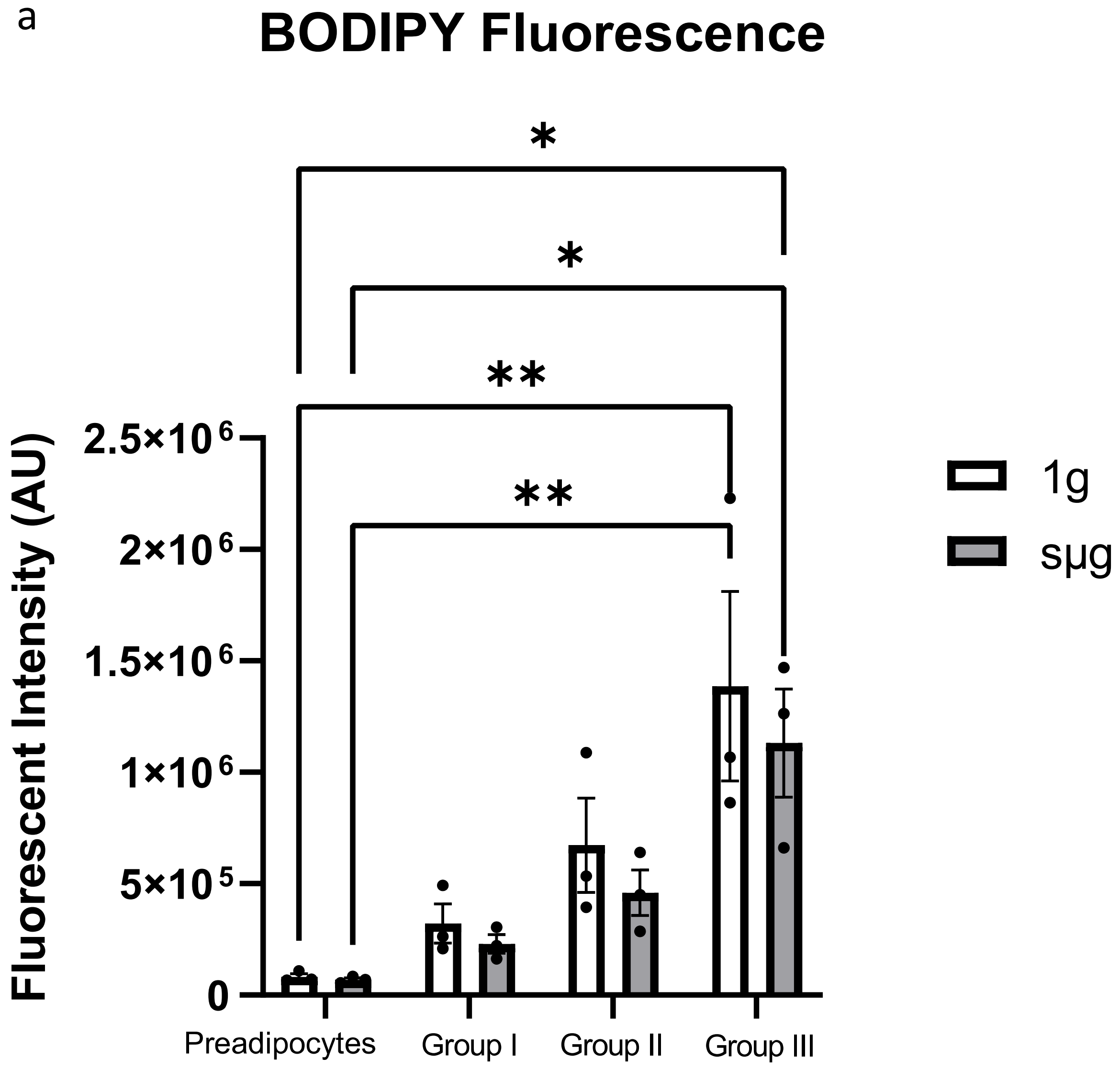

**Supplementary Figure S6. BODIPY fluorescent intensity in each subpopulation.** (a) Fluorescent intensity of BODIPY in each subpopulation for 1g and sμg conditions. Data are presented as means ± SEM. Comparisons between groups and statistical analysis were performed using two-way ANOVA with Tukey post hoc test. (*$p < 0.05$, **$p < 0.01$).

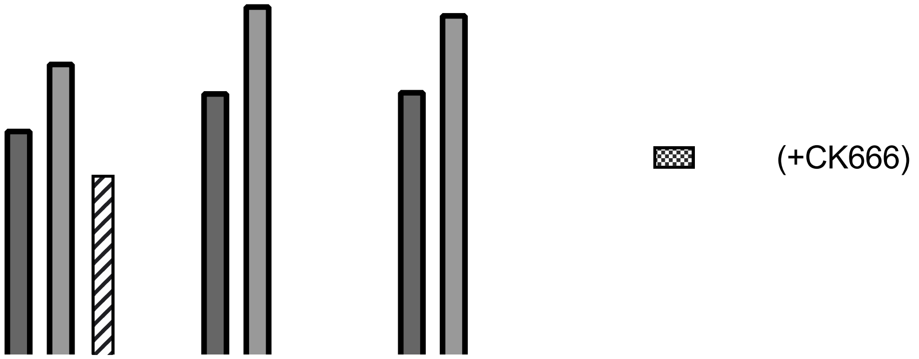

**Supplementary Figure S7. CK-666 inhibition of ARP2/3 remodeling did not affect insulin-stimulated glucose uptake in 1g condition.** (a) Insulin-stimulated glucose uptake was not significantly affected in 1g after exposure to CK-666 (n=3 biological replicates). Significant differences with CK-666 were only present in sμg condition. Data are presented as means ± SEM. Comparisons between groups and statistical analysis were performed using two-way ANOVA with Tukey post hoc test (*p<0.05, **p < 0.01, ****p<0.0001).

a

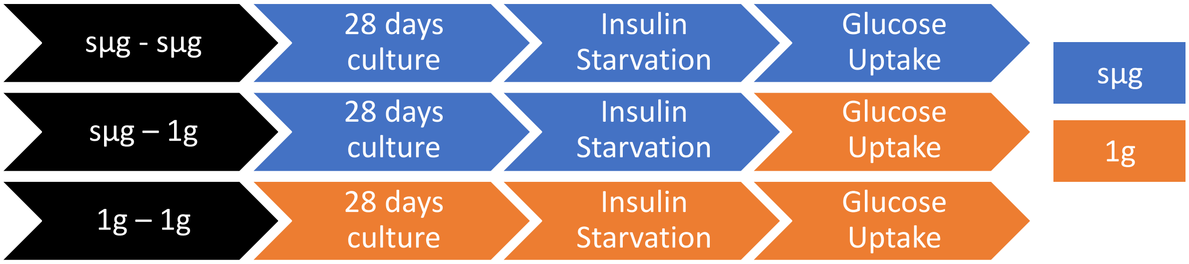

b

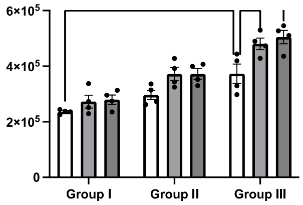

**Supplementary Figure S8. Insulin-stimulated glucose uptake in sμg condition was not reversed after a short-term exposure to 1g condition.** (a) Experimental workflow for performing insulin-stimulated glucose uptake in 3 swapped conditions. (b) Insulin-stimulated glucose uptake in sμg –1g conditions were similar to insulin-stimulated glucose uptake in sμg–sμg conditions (n=4 biological replicates). The sμg phenotype did not appear to be reversed after a short-term exposure to 1g conditions during insulin-stimulated glucose uptake (30 minutes). Comparisons between groups and statistical analysis were performed using two-way ANOVA with Tukey post hoc test (*p<0.05, **p < 0.01, ****p<0.0001).

a

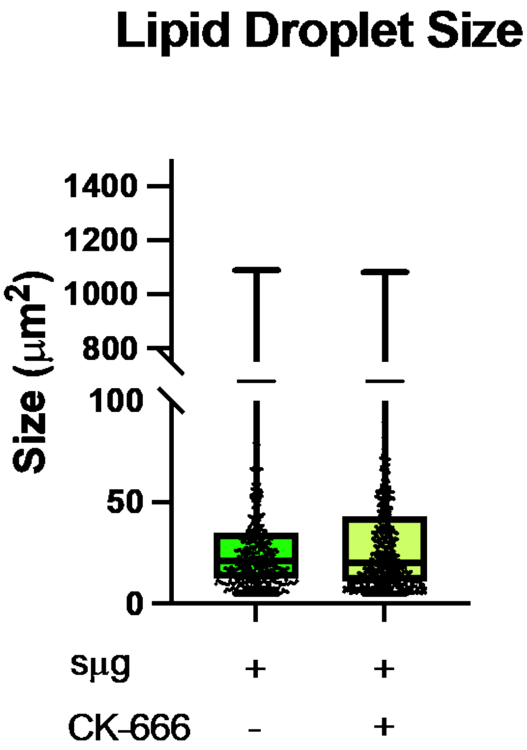

**Supplementary Figure S9. Lipid droplet size after exposure to CK-666.** (a) Lipid droplet size did not change in sμg in the presence of CK-666 after insulin-stimulated glucose uptake (n=3 biological replicates, 26 adipocytes total for lipid droplet sizes, Box and Whiskers graphs are presenting min to max data with their median). Data are presented as means ± SEM. Comparisons between groups and statistical analysis were performed using Mann–Whitney test.

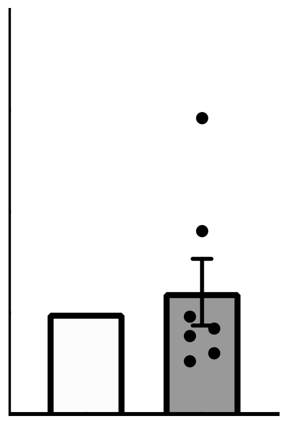

**Supplementary Figure S10. Adipocyte browning related genes.** (a) Among browning related genes, PPARG coactivator 1 alpha (*PGC-1α*) was significantly upregulated in sµg (n=6-7 biological replicates). Data are presented as means ± SEM. Comparisons between groups and statistical analysis were performed using an unpaired t-test with two-tailed p-values (*p < 0.05).

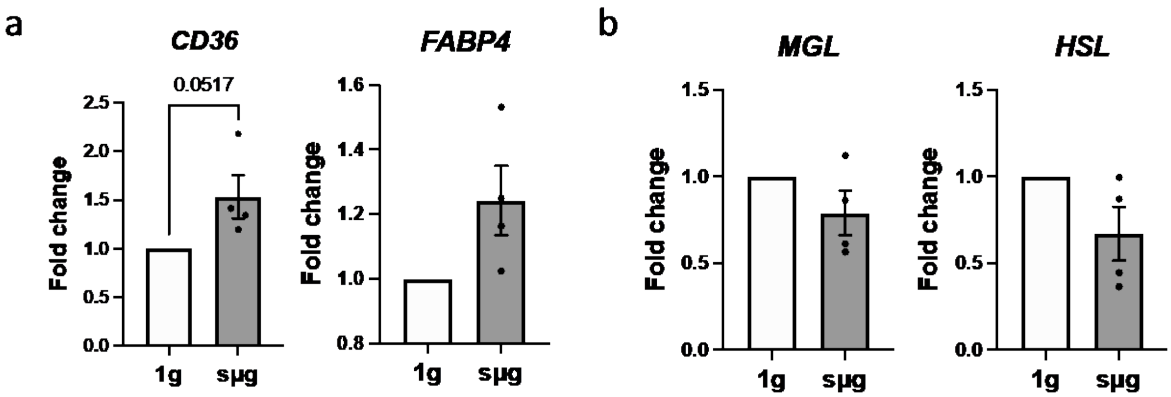

**Supplementary Figure S11. Additional lipogenesis and lipolysis related gene expression.** (a) Gene expression for lipogenesis, cluster of differentiation (*CD36*), fatty acid binding protein 4 (*FABP4*), and (b) lipolysis related genes, monoglyceride lipase (*MGL*) and hormone sensitive lipase (*HSL*) (n=4 biological replicates). Data are presented as means ± SEM. Comparisons between groups and statistical analysis were performed using an unpaired t-test with two-tailed p-values.

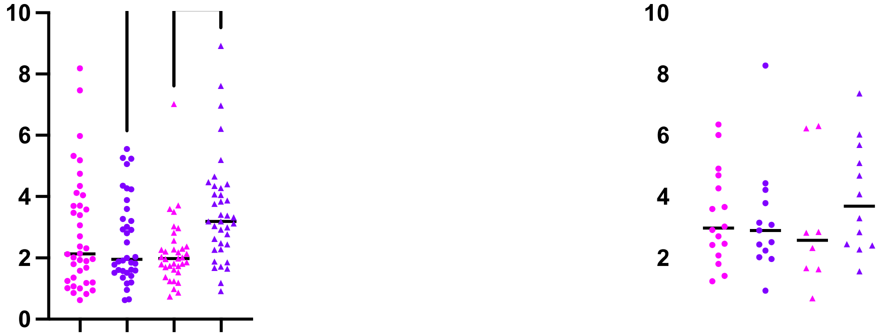

**Supplementary Figure S12. GLUT4 membrane to cytoplasm ratio was significantly upregulated in sµg for Group II.** GLUT4 membrane to cytoplasm ratio for (a) Group II (n=3 biological replicates, 10-14 cells were quantified for each replicate), and (b) Group III (n=3 biological replicates, 2-6 cells were quantified for each biological replicate). Data are presented as means ± SEM. Comparisons between groups and statistical analysis were performed using two-way ANOVA or Kruskal-Wallis test with Tukey post hoc test or Dunn's multiple comparisons test (*p < 0.05, **p < 0.01).

**Supplementary Table 1.** Primer sequences.

| Gene | Forward primer sequences | Reverse primer sequences |
|---|---|---|
| ***PPARG*** | ACCAAAGTGCAATCAAAGTGGA | ATGAGGGAGTTGGAAGGCTCT |
| ***ADIPOQ*** | GGCTTTCCGGGAATCCAAGG | TGGGGATAGTAACGTAAGTCTCC |
| ***LEP*** | TGCCTTCCAGAAACGTGATCC | CTCTGTGGAGTAGCCTGAAGC |
| ***RHOA*** | TGCCATCCGGAAGAAACTGG | GCCAACTCTACCTGCTTTCCA |
| ***ROCK1*** | TACTGACAGGGAAGTGAGGT | AATCGGGTACAACTGGTGCT |
| ***DGAT*** | TCTGGGAGATGGGGCACTG | GCTGCTTTTCCACCTTGGAC |
| ***PLIN-1*** | CTGTGAGACTGAGGTGGCG | ATTCTCCTGCTCAGGGAGGT |
| ***ATGL*** | GTCAGACGGCGAGAATGTCA | CTGCAGACATTGGCCTGGAT |
| ***CD36*** | GGACGCTGAGGACAACACAG | AGTTGTCAGCCTCTGTTCCAA |
| ***HSL*** | GACTGGCAACCTGAACCACA | TCGATTCTGGCTGGGCTATG |
| ***MGL*** | CCAGCATGCCAGAGGAAAGT | TGTCCGTCTGCATTGACCAG |
| ***UCP-1*** | GTGCCCAACTGTGCAATGAA | GAAGGAAGGTACCAACCCCT |
| ***PGC-1*** | CTCTCAGTAAGGGGCTGGTTG | CAACCAGAGCAGCACACTCG |
| ***FABP4*** | AAACTGGTGGTGGAATGCGT | GCGAACTTCAGTCCAGGTCA |
| ***HPRT-1*** | CTGGCGTCGTGATTAGTGAT | TCTCGAGCAAGACGTTCAGT |

**Supplementary Table 2.** Unpaired t-test statistical analysis of *PPARG* gene expression data in Figure 2.

| Table Analyzed | PPARG |
|---|---|
| Column B | sµg |
| vs. | vs. |
| Column A | 1g |
| Unpaired t test | |
| P value | 0.1998 |
| P value summary | ns |
| Significantly different (P < 0.05)? | No |
| One- or two-tailed P value? | Two-tailed |
| t, df | t=1.357, df=12 |
| How big is the difference? | |
| Mean of column A | 1.000 |
| Mean of column B | 1.537 |
| Difference between means (B - A) ± SEM | 0.5371 ± 0.3959 |
| 95% confidence interval | -0.3254 to 1.400 |
| R squared (eta squared) | 0.1330 |
| F test to compare variances | |
| F, DFn, Dfd | Infinity, 6, 6 |
| P value | <0.0001 |
| P value summary | **** |
| Significantly different (P < 0.05)? | Yes |
| Data analyzed | |
| Sample size, column A | 7 |
| Sample size, column B | 7 |

**Supplementary Table 3.** Unpaired t-test statistical analysis of *ADIPOQ* gene expression data in Figure 2.

| Table Analyzed | ADIPOQ |
|---|---|
| Column B | sµg |
| vs. | vs. |
| Column A | 1g |
| Unpaired t test | |
| P value | 0.3426 |
| P value summary | ns |
| Significantly different (P < 0.05)? | No |
| One- or two-tailed P value? | Two-tailed |
| t, df | t=0.9823, df=14 |
| How big is the difference? | |
| Mean of column A | 1.000 |
| Mean of column B | 1.363 |
| Difference between means (B - A) ± SEM | 0.3625 ± 0.3690 |
| 95% confidence interval | -0.4290 to 1.154 |
| R squared (eta squared) | 0.06447 |
| F test to compare variances | |
| F, DFn, Dfd | Infinity, 7, 7 |
| P value | <0.0001 |
| P value summary | **** |
| Significantly different (P < 0.05)? | Yes |
| Data analyzed | |
| Sample size, column A | 8 |
| Sample size, column B | 8 |

**Supplementary Table 4.** Unpaired t-test statistical analysis of *LEP* gene expression data in Figure 2.

| Table Analyzed | LEP |
|---|---|
| Column B | sµg |
| vs. | vs. |
| Column A | 1g |
| Unpaired t test with Welch's correction | |
| P value | <0.0001 |
| P value summary | **** |
| Significantly different (P < 0.05)? | Yes |
| One- or two-tailed P value? | Two-tailed |
| Welch-corrected t, df | t=16.85, df=7.000 |
| How big is the difference? | |
| Mean of column A | 1.000 |
| Mean of column B | 0.1663 |
| Difference between means (B - A) ± SEM | -0.8338 ± 0.04950 |
| 95% confidence interval | -0.9508 to -0.7167 |
| R squared (eta squared) | 0.9759 |
| F test to compare variances | |
| F, DFn, Dfd | Infinity, 7, 7 |
| P value | <0.0001 |
| P value summary | **** |
| Significantly different (P < 0.05)? | Yes |
| Data analyzed | |
| Sample size, column A | 8 |
| Sample size, column B | 8 |

**Supplementary Table 5.** Normality test result for Area data in Figure 2.

| D'Agostino & Pearson test | | |
|---|---|---|
| K2 | 2.733 | 13.42 |
| P value | 0.2550 | 0.0012 |
| Passed normality test (alpha=0.05)? | Yes | No |
| P value summary | ns | ** |

**Supplementary Table 6.** Mann-Whitney test statistical analysis of Area data in Figure 2.

| Table Analyzed | Area |
|---|---|
| Column B | sμg |
| vs. | vs. |
| Column A | 1g |
| Mann Whitney test | |
| P value | 0.0001 |
| Exact or approximate P value? | Exact |
| P value summary | *** |
| Significantly different (P < 0.05)? | Yes |
| One- or two-tailed P value? | Two-tailed |
| Sum of ranks in column A,B | 6765 , 9707 |
| Mann-Whitney U | 2760 |
| Difference between medians | |
| Median of column A | 701.7, n=89 |
| Median of column B | 888.8, n=92 |
| Difference: Actual | 187.1 |
| Difference: Hodges-Lehmann | 191.9 |

**Supplementary Table 7.** Normality test result for Perimeter data in Figure 2.

| D'Agostino & Pearson test | | |
|---|---|---|
| K2 | 0.6244 | 26.51 |
| P value | 0.7318 | <0.0001 |
| Passed normality test (alpha=0.05)? | Yes | No |
| P value summary | ns | **** |

**Supplementary Table 8.** Mann-Whitney test statistical analysis of Perimeter data in Figure 2.

| Table Analyzed | Perimeter |
|---|---|
| Column B | sμg |
| vs. | vs. |
| Column A | 1g |
| Mann Whitney test | |
| P value | 0.0003 |
| Exact or approximate P value? | Exact |
| P value summary | *** |
| Significantly different (P < 0.05)? | Yes |
| One- or two-tailed P value? | Two-tailed |
| Sum of ranks in column A,B | 6833 , 9639 |
| Mann-Whitney U | 2828 |
| Difference between medians | |
| Median of column A | 105.1, n=89 |
| Median of column B | 120.2, n=92 |
| Difference: Actual | 15.18 |
| Difference: Hodges-Lehmann | 13.49 |

**Supplementary Table 9.** Normality test result for Lipid Droplet Size data in Figure 2.

| D'Agostino & Pearson test | | |
|---|---|---|
| K2 | 895.9 | 613.7 |
| P value | <0.0001 | <0.0001 |
| Passed normality test (alpha=0.05)? | No | No |
| P value summary | **** | **** |

**Supplementary Table 10.** Mann-Whitney test statistical analysis of Lipid Droplet Size data in Figure 2.

| Table Analyzed | Lipid Droplet Size |
|---|---|
| Column B | sμg |
| vs. | vs. |
| Column A | 1g |
| Mann Whitney test | |
| P value | 0.9785 |
| Exact or approximate P value? | Approximate |
| P value summary | ns |
| Significantly different (P < 0.05)? | No |
| One- or two-tailed P value? | Two-tailed |
| Sum of ranks in column A,B | 836153 , 828247 |
| Mann-Whitney U | 415561 |
| Difference between medians | |
| Median of column A | 16.63, n=916 |
| Median of column B | 17.14, n=908 |
| Difference: Actual | 0.5150 |
| Difference: Hodges-Lehmann | 0.000 |

**Supplementary Table 11.** Two-way ANOVA test statistical analysis of Basal Lipolysis data in Figure 2.

| Table Analyzed | Basal Lipolysis | | | | |
|---|---|---|---|---|---|
| Two-way ANOVA | Ordinary | | | | |
| Alpha | 0.05 | | | | |
| Source of Variation | % of total variation | P value | P value summary | Significant? | |
| Interaction | 11.22 | 0.0012 | ** | Yes | |
| Row Factor | 40.07 | <0.0001 | **** | Yes | |
| Column Factor | 30.90 | <0.0001 | **** | Yes | |
| ANOVA table | SS | DF | MS | F (DFn, DFd) | P value |
| Interaction | 0.03179 | 3 | 0.01060 | F (3, 32) = 6.715 | P=0.0012 |
| Row Factor | 0.1136 | 3 | 0.03786 | F (3, 32) = 23.99 | P<0.0001 |
| Column Factor | 0.08761 | 1 | 0.08761 | F (1, 32) = 55.51 | P<0.0001 |
| Residual | 0.05051 | 32 | 0.001578 | | |
| Difference between column means | | | | | |
| Mean of 1g | 0.2813 | | | | |
| Mean of sμg | 0.3749 | | | | |
| Difference between means | -0.09360 | | | | |
| SE of difference | 0.01256 | | | | |
| 95% CI of difference | -0.1192 to -0.06801 | | | | |
| Data summary | | | | | |
| Number of columns (Column Factor) | 2 | | | | |
| Number of rows (Row Factor) | 4 | | | | |
| Number of values | 40 | | | | |

**Supplementary Table 12.** Multiple comparisons results for Two-way ANOVA test statistical analysis of Basal Lipolysis data in Figure 2.

Compare cell means regardless of rows and columns

| | |
|---|---|
| Number of families | 1 |
| Number of comparisons per family | 28 |
| Alpha | 0.05 |

| Tukey's multiple comparisons test | Mean Diff. | 95.00% CI of diff. | Below threshold? | Summary | Adjusted P Value |
|---|---|---|---|---|---|
| Row 1:1g vs. Row 1:spg | -0.005658 | -0.08705 to 0.07573 | No | ns | >0.9999 |
| Row 1:1g vs. Row 2:1g | -0.01775 | -0.09914 to 0.06364 | No | ns | 0.9962 |
| Row 1:1g vs. Row 2:spg | -0.1006 | -0.1820 to -0.01923 | Yes | ** | 0.0074 |
| Row 1:1g vs. Row 3:1g | -0.04476 | -0.1261 to 0.03664 | No | ns | 0.6365 |
| Row 1:1g vs. Row 3:spg | -0.1871 | -0.2685 to -0.1057 | Yes | **** | <0.0001 |
| Row 1:1g vs. Row 4:1g | -0.06890 | -0.1503 to 0.01249 | No | ns | 0.1467 |
| Row 1:1g vs. Row 4:spg | -0.2124 | -0.2938 to -0.1310 | Yes | **** | <0.0001 |
| Row 1:spg vs. Row 2:1g | -0.01209 | -0.09348 to 0.06930 | No | ns | 0.9997 |
| Row 1:spg vs. Row 2:spg | -0.09497 | -0.1764 to -0.01357 | Yes | * | 0.0133 |
| Row 1:spg vs. Row 3:1g | -0.03910 | -0.1205 to 0.04229 | No | ns | 0.7716 |
| Row 1:spg vs. Row 3:spg | -0.1815 | -0.2629 to -0.1001 | Yes | **** | <0.0001 |
| Row 1:spg vs. Row 4:1g | -0.06324 | -0.1446 to 0.01815 | No | ns | 0.2247 |
| Row 1:spg vs. Row 4:spg | -0.2067 | -0.2881 to -0.1253 | Yes | **** | <0.0001 |
| Row 2:1g vs. Row 2:spg | -0.08288 | -0.1643 to -0.001485 | Yes | * | 0.0435 |
| Row 2:1g vs. Row 3:1g | -0.02701 | -0.1084 to 0.05438 | No | ns | 0.9575 |
| Row 2:1g vs. Row 3:spg | -0.1694 | -0.2508 to -0.08800 | Yes | **** | <0.0001 |
| Row 2:1g vs. Row 4:1g | -0.05115 | -0.1325 to 0.03024 | No | ns | 0.4752 |
| Row 2:1g vs. Row 4:spg | -0.1946 | -0.2760 to -0.1133 | Yes | **** | <0.0001 |
| Row 2:spg vs. Row 3:1g | 0.05587 | -0.02552 to 0.1373 | No | ns | 0.3652 |
| Row 2:spg vs. Row 3:spg | -0.08651 | -0.1679 to -0.005122 | Yes | * | 0.0308 |
| Row 2:spg vs. Row 4:1g | 0.03172 | -0.04967 to 0.1131 | No | ns | 0.9058 |
| Row 2:spg vs. Row 4:spg | -0.1118 | -0.1932 to -0.03038 | Yes | ** | 0.0022 |
| Row 3:1g vs. Row 3:spg | -0.1424 | -0.2238 to -0.06099 | Yes | **** | <0.0001 |
| Row 3:1g vs. Row 4:1g | -0.02415 | -0.1055 to 0.05725 | No | ns | 0.9768 |
| Row 3:1g vs. Row 4:spg | -0.1676 | -0.2490 to -0.08625 | Yes | **** | <0.0001 |
| Row 3:spg vs. Row 4:1g | 0.1182 | 0.03684 to 0.1996 | Yes | ** | 0.0011 |
| Row 3:spg vs. Row 4:spg | -0.02526 | -0.1066 to 0.05613 | No | ns | 0.9703 |
| Row 4:1g vs. Row 4:spg | -0.1435 | -0.2249 to -0.06210 | Yes | **** | <0.0001 |

| Test details | Mean 1 | Mean 2 | Mean Diff. | SE of diff. | N1 | N2 | q | DF |
|---|---|---|---|---|---|---|---|---|
| Row 1:1g vs. Row 1:spg | 0.2485 | 0.2541 | -0.005658 | 0.02513 | 5 | 5 | 0.3184 | 32.00 |
| Row 1:1g vs. Row 2:1g | 0.2485 | 0.2662 | -0.01775 | 0.02513 | 5 | 5 | 0.9989 | 32.00 |
| Row 1:1g vs. Row 2:spg | 0.2485 | 0.3491 | -0.1006 | 0.02513 | 5 | 5 | 5.664 | 32.00 |
| Row 1:1g vs. Row 3:1g | 0.2485 | 0.2932 | -0.04476 | 0.02513 | 5 | 5 | 2.519 | 32.00 |
| Row 1:1g vs. Row 3:spg | 0.2485 | 0.4356 | -0.1871 | 0.02513 | 5 | 5 | 10.53 | 32.00 |
| Row 1:1g vs. Row 4:1g | 0.2485 | 0.3174 | -0.06890 | 0.02513 | 5 | 5 | 3.878 | 32.00 |
| Row 1:1g vs. Row 4:spg | 0.2485 | 0.4609 | -0.2124 | 0.02513 | 5 | 5 | 11.95 | 32.00 |
| Row 1:spg vs. Row 2:1g | 0.2541 | 0.2662 | -0.01209 | 0.02513 | 5 | 5 | 0.6805 | 32.00 |
| Row 1:spg vs. Row 2:spg | 0.2541 | 0.3491 | -0.09497 | 0.02513 | 5 | 5 | 5.345 | 32.00 |
| Row 1:spg vs. Row 3:1g | 0.2541 | 0.2932 | -0.03910 | 0.02513 | 5 | 5 | 2.201 | 32.00 |
| Row 1:spg vs. Row 3:spg | 0.2541 | 0.4356 | -0.1815 | 0.02513 | 5 | 5 | 10.21 | 32.00 |
| Row 1:spg vs. Row 4:1g | 0.2541 | 0.3174 | -0.06324 | 0.02513 | 5 | 5 | 3.560 | 32.00 |
| Row 1:spg vs. Row 4:spg | 0.2541 | 0.4609 | -0.2067 | 0.02513 | 5 | 5 | 11.64 | 32.00 |
| Row 2:1g vs. Row 2:spg | 0.2662 | 0.3491 | -0.08288 | 0.02513 | 5 | 5 | 4.665 | 32.00 |
| Row 2:1g vs. Row 3:1g | 0.2662 | 0.2932 | -0.02701 | 0.02513 | 5 | 5 | 1.520 | 32.00 |
| Row 2:1g vs. Row 3:spg | 0.2662 | 0.4356 | -0.1694 | 0.02513 | 5 | 5 | 9.534 | 32.00 |
| Row 2:1g vs. Row 4:1g | 0.2662 | 0.3174 | -0.05115 | 0.02513 | 5 | 5 | 2.879 | 32.00 |
| Row 2:1g vs. Row 4:spg | 0.2662 | 0.4609 | -0.1946 | 0.02513 | 5 | 5 | 10.96 | 32.00 |
| Row 2:spg vs. Row 3:1g | 0.3491 | 0.2932 | 0.05587 | 0.02513 | 5 | 5 | 3.145 | 32.00 |
| Row 2:spg vs. Row 3:spg | 0.3491 | 0.4356 | -0.08651 | 0.02513 | 5 | 5 | 4.869 | 32.00 |
| Row 2:spg vs. Row 4:1g | 0.3491 | 0.3174 | 0.03172 | 0.02513 | 5 | 5 | 1.785 | 32.00 |
| Row 2:spg vs. Row 4:spg | 0.3491 | 0.4609 | -0.1118 | 0.02513 | 5 | 5 | 6.291 | 32.00 |
| Row 3:1g vs. Row 3:spg | 0.2932 | 0.4356 | -0.1424 | 0.02513 | 5 | 5 | 8.014 | 32.00 |
| Row 3:1g vs. Row 4:1g | 0.2932 | 0.3174 | -0.02415 | 0.02513 | 5 | 5 | 1.359 | 32.00 |
| Row 3:1g vs. Row 4:spg | 0.2932 | 0.4609 | -0.1676 | 0.02513 | 5 | 5 | 9.435 | 32.00 |
| Row 3:spg vs. Row 4:1g | 0.4356 | 0.3174 | 0.1182 | 0.02513 | 5 | 5 | 6.655 | 32.00 |
| Row 3:spg vs. Row 4:spg | 0.4356 | 0.4609 | -0.02526 | 0.02513 | 5 | 5 | 1.422 | 32.00 |
| Row 4:1g vs. Row 4:spg | 0.3174 | 0.4609 | -0.1435 | 0.02513 | 5 | 5 | 8.076 | 32.00 |

**Supplementary Table 13.** Unpaired t-test statistical analysis of *ROCK1* gene expression data in Figure 3.

| Table Analyzed | ROCK1 |
|---|---|
| Column B | sμg |
| vs. | vs. |
| Column A | 1g |
| Unpaired t test | |
| P value | 0.0195 |
| P value summary | * |
| Significantly different (P < 0.05)? | Yes |
| One- or two-tailed P value? | Two-tailed |
| t, df | t=2.914, df=8 |
| How big is the difference? | |
| Mean of column A | 1.000 |
| Mean of column B | 1.434 |
| Difference between means (B - A) ± SEM | 0.4340 ± 0.1489 |
| 95% confidence interval | 0.09060 to 0.7774 |
| R squared (eta squared) | 0.5150 |
| F test to compare variances | |
| F, DFn, Dfd | Infinity, 4, 4 |
| P value | <0.0001 |
| P value summary | **** |
| Significantly different (P < 0.05)? | Yes |
| Data analyzed | |
| Sample size, column A | 5 |
| Sample size, column B | 5 |

**Supplementary Table 14.** Unpaired t-test statistical analysis of *RHOA* gene expression data in Figure 3.

| Table Analyzed | RHOA |
|---|---|
| Column B | sμg |
| vs. | vs. |
| Column A | 1g |
| Unpaired t test | |
| P value | 0.0137 |
| P value summary | * |
| Significantly different (P < 0.05)? | Yes |
| One- or two-tailed P value? | Two-tailed |
| t, df | t=3.146, df=8 |
| How big is the difference? | |
| Mean of column A | 1.000 |
| Mean of column B | 1.938 |
| Difference between means (B - A) ± SEM | 0.9380 ± 0.2982 |
| 95% confidence interval | 0.2504 to 1.626 |
| R squared (eta squared) | 0.5530 |
| F test to compare variances | |
| F, DFn, Dfd | Infinity, 4, 4 |
| P value | <0.0001 |
| P value summary | **** |
| Significantly different (P < 0.05)? | Yes |
| Data analyzed | |
| Sample size, column A | 5 |
| Sample size, column B | 5 |

**Supplementary Table 15.** Unpaired t-test statistical analysis of Total RhoA protein expression data in Figure 3.

| Table Analyzed | Total RhoA |
|---|---|
| Column B | sµg |
| vs. | vs. |
| Column A | 1g |
| Unpaired t test | |
| P value | 0.9899 |
| P value summary | ns |
| Significantly different (P < 0.05)? | No |
| One- or two-tailed P value? | Two-tailed |
| t, df | t=0.01341, df=4 |
| How big is the difference? | |
| Mean of column A | 1.000 |
| Mean of column B | 1.003 |
| Difference between means (B - A) ± SEM | 0.003333 ± 0.2485 |
| 95% confidence interval | -0.6866 to 0.6932 |
| R squared (eta squared) | 4.499e-005 |
| F test to compare variances | |
| F, DFn, Dfd | Infinity, 2, 2 |
| P value | <0.0001 |
| P value summary | **** |
| Significantly different (P < 0.05)? | Yes |
| Data analyzed | |
| Sample size, column A | 3 |
| Sample size, column B | 3 |

**Supplementary Table 16.** Unpaired t-test statistical analysis of ARP3 protein expression data in Figure 3.

| Table Analyzed | ARP3 |
|---|---|
| Column B | sµg |
| vs. | vs. |
| Column A | 1g |
| Unpaired t test | |
| P value | 0.0245 |
| P value summary | * |
| Significantly different (P < 0.05)? | Yes |
| One- or two-tailed P value? | Two-tailed |
| t, df | t=3.516, df=4 |
| How big is the difference? | |
| Mean of column A | 1.000 |
| Mean of column B | 1.903 |
| Difference between means (B - A) ± SEM | 0.9033 ± 0.2569 |
| 95% confidence interval | 0.1900 to 1.617 |
| R squared (eta squared) | 0.7555 |
| F test to compare variances | |
| F, DFn, Dfd | Infinity, 2, 2 |
| P value | <0.0001 |
| P value summary | **** |
| Significantly different (P < 0.05)? | Yes |
| Data analyzed | |
| Sample size, column A | 3 |
| Sample size, column B | 3 |

**Supplementary Table 17.** Unpaired t-test statistical analysis of pCofilin/Cofilin protein expression data in Figure 3.

| Table Analyzed | pCofilin/Cofilin |
|---|---|
| Column B | sμg |
| vs. | vs. |
| Column A | 1g |
| Unpaired t test | |
| P value | 0.0285 |
| P value summary | * |
| Significantly different (P < 0.05)? | Yes |
| One- or two-tailed P value? | Two-tailed |
| t, df | t=2.558, df=10 |
| How big is the difference? | |
| Mean of column A | 1.000 |
| Mean of column B | 1.923 |
| Difference between means (B - A) ± SEM | 0.9233 ± 0.3609 |
| 95% confidence interval | 0.1191 to 1.728 |
| R squared (eta squared) | 0.3956 |
| F test to compare variances | |
| F, DFn, Dfd | Infinity, 5, 5 |
| P value | <0.0001 |
| P value summary | **** |
| Significantly different (P < 0.05)? | Yes |
| Data analyzed | |
| Sample size, column A | 6 |
| Sample size, column B | 6 |

**Supplementary Table 18.** Unpaired t-test statistical analysis of ARP2 protein expression data in Figure 3.

| Table Analyzed | ARP2 |
|---|---|
| Column B | sμg |
| vs. | vs. |
| Column A | 1g |
| Unpaired t test | |
| P value | 0.9495 |
| P value summary | ns |
| Significantly different (P < 0.05)? | No |
| One- or two-tailed P value? | Two-tailed |
| t, df | t=0.06744, df=4 |
| How big is the difference? | |
| Mean of column A | 1.000 |
| Mean of column B | 0.9733 |
| Difference between means (B - A) ± SEM | -0.02667 ± 0.3954 |
| 95% confidence interval | -1.124 to 1.071 |
| R squared (eta squared) | 0.001136 |
| F test to compare variances | |
| F, DFn, Dfd | Infinity, 2, 2 |
| P value | <0.0001 |
| P value summary | **** |
| Significantly different (P < 0.05)? | Yes |
| Data analyzed | |
| Sample size, column A | 3 |
| Sample size, column B | 3 |

**Supplementary Table 19.** Two-way ANOVA test statistical analysis of Adipocyte SSCA-A Subpopulations data in Figure 4.

| Table Analyzed | Adipocyte SSC-A Subpopulations | | | | |
|---|---|---|---|---|---|
| Two-way ANOVA | Ordinary | | | | |
| Alpha | 0.05 | | | | |
| Source of Variation | % of total variation | P value | P value summary | Significant? | |
| Interaction | 13.22 | 0.0031 | ** | Yes | |
| Row Factor | 72.13 | <0.0001 | **** | Yes | |
| Column Factor | 0.000 | >0.9999 | ns | No | |
| ANOVA table | SS | DF | MS | F (DFn, DFd) | P value |
| Interaction | 519.9 | 2 | 259.9 | F (2, 18) = 8.124 | P=0.0031 |
| Row Factor | 2836 | 2 | 1418 | F (2, 18) = 44.32 | P<0.0001 |
| Column Factor | 0.000 | 1 | 0.000 | F (1, 18) = 0.000 | P>0.9999 |
| Residual | 576.0 | 18 | 32.00 | | |
| Difference between column means | | | | | |
| Mean of 1g | 33.33 | | | | |
| Mean of sµg | 33.33 | | | | |
| Difference between means | 0.000 | | | | |
| SE of difference | 2.309 | | | | |
| 95% CI of difference | -4.852 to 4.852 | | | | |
| Data summary | | | | | |
| Number of columns (Column Factor) | 2 | | | | |
| Number of rows (Row Factor) | 3 | | | | |
| Number of values | 24 | | | | |

**Supplementary Table 20.** Multiple comparisons results for Two-way ANOVA test statistical analysis of Adipocyte SSCA-A Subpopulations data in Figure 4.

**Compare cell means regardless of rows and columns**

| | |
|---|---|
| **Number of families** | **1** |
| **Number of comparisons per family** | **15** |
| **Alpha** | **0.05** |

| Tukey's multiple comparisons test | Mean Diff. | 95.00% CI of diff. | Below threshold? | Summary | Adjusted P Value |
|---|---|---|---|---|---|
| Group I:1g vs. Group I:sμg | 9.318 | -3.394 to 22.03 | No | ns | 0.2330 |
| Group I:1g vs. Group II:1g | 28.86 | 16.14 to 41.57 | Yes | **** | <0.0001 |
| Group I:1g vs. Group II:sμg | 32.25 | 19.54 to 44.96 | Yes | **** | <0.0001 |
| Group I:1g vs. Group III:1g | 29.34 | 16.63 to 42.05 | Yes | **** | <0.0001 |
| Group I:1g vs. Group III:sμg | 16.63 | 3.913 to 29.34 | Yes | ** | 0.0066 |
| Group I:sμg vs. Group II:1g | 19.54 | 6.826 to 32.25 | Yes | ** | 0.0014 |
| Group I:sμg vs. Group II:sμg | 22.93 | 10.22 to 35.64 | Yes | *** | 0.0002 |
| Group I:sμg vs. Group III:1g | 20.02 | 7.308 to 32.73 | Yes | ** | 0.0011 |
| Group I:sμg vs. Group III:sμg | 7.307 | -5.404 to 20.02 | No | ns | 0.4743 |
| Group II:1g vs. Group II:sμg | 3.395 | -9.317 to 16.11 | No | ns | 0.9538 |
| Group II:1g vs. Group III:1g | 0.4825 | -12.23 to 13.19 | No | ns | >0.9999 |
| Group II:1g vs. Group III:sμg | -12.23 | -24.94 to 0.4819 | No | ns | 0.0633 |
| Group II:sμg vs. Group III:1g | -2.913 | -15.62 to 9.799 | No | ns | 0.9757 |
| Group II:sμg vs. Group III:sμg | -15.63 | -28.34 to -2.913 | Yes | * | 0.0112 |
| Group III:1g vs. Group III:sμg | -12.71 | -25.42 to -0.0006389 | Yes | * | 0.0500 |

| Test details | Mean 1 | Mean 2 | Mean Diff. | SE of diff. | N1 | N2 | q | DF |
|---|---|---|---|---|---|---|---|---|
| Group I:1g vs. Group I:sμg | 52.73 | 43.41 | 9.318 | 4.000 | 4 | 4 | 3.294 | 18.00 |
| Group I:1g vs. Group II:1g | 52.73 | 23.88 | 28.86 | 4.000 | 4 | 4 | 10.20 | 18.00 |
| Group I:1g vs. Group II:sμg | 52.73 | 20.48 | 32.25 | 4.000 | 4 | 4 | 11.40 | 18.00 |
| Group I:1g vs. Group III:1g | 52.73 | 23.39 | 29.34 | 4.000 | 4 | 4 | 10.37 | 18.00 |
| Group I:1g vs. Group III:sμg | 52.73 | 36.11 | 16.63 | 4.000 | 4 | 4 | 5.878 | 18.00 |
| Group I:sμg vs. Group II:1g | 43.41 | 23.88 | 19.54 | 4.000 | 4 | 4 | 6.908 | 18.00 |
| Group I:sμg vs. Group II:sμg | 43.41 | 20.48 | 22.93 | 4.000 | 4 | 4 | 8.108 | 18.00 |
| Group I:sμg vs. Group III:1g | 43.41 | 23.39 | 20.02 | 4.000 | 4 | 4 | 7.078 | 18.00 |
| Group I:sμg vs. Group III:sμg | 43.41 | 36.11 | 7.307 | 4.000 | 4 | 4 | 2.584 | 18.00 |
| Group II:1g vs. Group II:sμg | 23.88 | 20.48 | 3.395 | 4.000 | 4 | 4 | 1.200 | 18.00 |
| Group II:1g vs. Group III:1g | 23.88 | 23.39 | 0.4825 | 4.000 | 4 | 4 | 0.1706 | 18.00 |
| Group II:1g vs. Group III:sμg | 23.88 | 36.11 | -12.23 | 4.000 | 4 | 4 | 4.324 | 18.00 |
| Group II:sμg vs. Group III:1g | 20.48 | 23.39 | -2.913 | 4.000 | 4 | 4 | 1.030 | 18.00 |
| Group II:sμg vs. Group III:sμg | 20.48 | 36.11 | -15.63 | 4.000 | 4 | 4 | 5.524 | 18.00 |
| Group III:1g vs. Group III:sμg | 23.39 | 36.11 | -12.71 | 4.000 | 4 | 4 | 4.495 | 18.00 |

**Supplementary Table 21.** Two-way ANOVA test statistical analysis of Glucose Uptake data in Figure 5.

| Table Analyzed | Glucose Uptake | | | | |
|---|---|---|---|---|---|
| Two-way ANOVA | Ordinary | | | | |
| Alpha | 0.05 | | | | |
| Source of Variation | % of total variation | P value | P value summary | Significant? | |
| Interaction | 3.740 | 0.1405 | ns | No | |
| Row Factor | 61.33 | <0.0001 | **** | Yes | |
| Column Factor | 19.58 | 0.0001 | *** | Yes | |
| ANOVA table | SS | DF | MS | F (DFn, DFd) | P value |
| Interaction | 8044768135 | 2 | 4022384067 | F (2, 18) = 2.193 | P=0.1405 |
| Row Factor | 131902606105 | 2 | 65951303053 | F (2, 18) = 35.95 | P<0.0001 |
| Column Factor | 42110450413 | 1 | 42110450413 | F (1, 18) = 22.96 | P=0.0001 |
| Residual | 33017275575 | 18 | 1834293088 | | |
| Difference between column means | | | | | |
| Mean of 1g | 301633 | | | | |
| Mean of sµg | 385409 | | | | |
| Difference between means | -83776 | | | | |
| SE of difference | 17485 | | | | |
| 95% CI of difference | -120510 to -47042 | | | | |
| Data summary | | | | | |
| Number of columns (Column Factor) | 2 | | | | |
| Number of rows (Row Factor) | 3 | | | | |
| Number of values | 24 | | | | |

**Supplementary Table 22.** Multiple comparisons results for Two-way ANOVA test statistical analysis of Glucose Uptake data in Figure 5.

Compare cell means regardless of rows and columns

| | |
|---|---|
| Number of families | 1 |
| Number of comparisons per family | 15 |
| Alpha | 0.05 |

| Tukey's multiple comparisons test | Mean Diff. | 95.00% CI of diff. | Below threshold? | Summary | Adjusted P Value |
|---|---|---|---|---|---|
| Group I:1g vs. Group I:sμg | -43656 | -139901 to 52589 | No | ns | 0.7028 |
| Group I:1g vs. Group II:1g | -60562 | -156807 to 35683 | No | ns | 0.3799 |
| Group I:1g vs. Group II:sμg | -136044 | -232289 to -39799 | Yes | ** | 0.0033 |
| Group I:1g vs. Group III:1g | -136608 | -232853 to -40363 | Yes | ** | 0.0031 |
| Group I:1g vs. Group III:sμg | -268798 | -365043 to -172553 | Yes | **** | <0.0001 |
| Group I:sμg vs. Group II:1g | -16906 | -113151 to 79338 | No | ns | 0.9926 |
| Group I:sμg vs. Group II:sμg | -92388 | -188633 to 3857 | No | ns | 0.0642 |
| Group I:sμg vs. Group III:1g | -92952 | -189197 to 3293 | No | ns | 0.0619 |
| Group I:sμg vs. Group III:sμg | -225142 | -321387 to -128897 | Yes | **** | <0.0001 |
| Group II:1g vs. Group II:sμg | -75482 | -171727 to 20763 | No | ns | 0.1781 |
| Group II:1g vs. Group III:1g | -76045 | -172290 to 20200 | No | ns | 0.1725 |
| Group II:1g vs. Group III:sμg | -208236 | -304481 to -111991 | Yes | **** | <0.0001 |
| Group II:sμg vs. Group III:1g | -563.7 | -96809 to 95681 | No | ns | >0.9999 |
| Group II:sμg vs. Group III:sμg | -132754 | -228999 to -36509 | Yes | ** | 0.0041 |
| Group III:1g vs. Group III:sμg | -132190 | -228435 to -35945 | Yes | ** | 0.0043 |

| Test details | Mean 1 | Mean 2 | Mean Diff. | SE of diff. | N1 | N2 | q | DF |
|---|---|---|---|---|---|---|---|---|
| Group I:1g vs. Group I:sμg | 235909 | 279565 | -43656 | 30284 | 4 | 4 | 2.039 | 18.00 |
| Group I:1g vs. Group II:1g | 235909 | 296472 | -60562 | 30284 | 4 | 4 | 2.828 | 18.00 |
| Group I:1g vs. Group II:sμg | 235909 | 371953 | -136044 | 30284 | 4 | 4 | 6.353 | 18.00 |
| Group I:1g vs. Group III:1g | 235909 | 372517 | -136608 | 30284 | 4 | 4 | 6.379 | 18.00 |
| Group I:1g vs. Group III:sμg | 235909 | 504707 | -268798 | 30284 | 4 | 4 | 12.55 | 18.00 |
| Group I:sμg vs. Group II:1g | 279565 | 296472 | -16906 | 30284 | 4 | 4 | 0.7895 | 18.00 |
| Group I:sμg vs. Group II:sμg | 279565 | 371953 | -92388 | 30284 | 4 | 4 | 4.314 | 18.00 |
| Group I:sμg vs. Group III:1g | 279565 | 372517 | -92952 | 30284 | 4 | 4 | 4.341 | 18.00 |
| Group I:sμg vs. Group III:sμg | 279565 | 504707 | -225142 | 30284 | 4 | 4 | 10.51 | 18.00 |
| Group II:1g vs. Group II:sμg | 296472 | 371953 | -75482 | 30284 | 4 | 4 | 3.525 | 18.00 |
| Group II:1g vs. Group III:1g | 296472 | 372517 | -76045 | 30284 | 4 | 4 | 3.551 | 18.00 |
| Group II:1g vs. Group III:sμg | 296472 | 504707 | -208236 | 30284 | 4 | 4 | 9.724 | 18.00 |
| Group II:sμg vs. Group III:1g | 371953 | 372517 | -563.7 | 30284 | 4 | 4 | 0.02632 | 18.00 |
| Group II:sμg vs. Group III:sμg | 371953 | 504707 | -132754 | 30284 | 4 | 4 | 6.199 | 18.00 |
| Group III:1g vs. Group III:sμg | 372517 | 504707 | -132190 | 30284 | 4 | 4 | 6.173 | 18.00 |

**Supplementary Table 23.** Normality test result for GLUT4 Fluorescent Intensity (Groups II,III) data in Figure 5.

| D'Agostino & Pearson test | | | | |
|---|---|---|---|---|
| K2 | 3.267 | 7.312 | 24.26 | 21.16 |
| P value | 0.1953 | 0.0258 | <0.0001 | <0.0001 |
| Passed normality test (alpha=0.05)? | Yes | No | No | No |
| P value summary | ns | * | **** | **** |

**Supplementary Table 24.** ANOVA results for Kruskal-Wallis test for GLUT4 Fluorescent Intensity (Groups II,III) data in Figure 5.

| Table Analyzed | GLUT4 (Groups II,III) |
|---|---|
| Kruskal-Wallis test | |
| P value | <0.0001 |
| Exact or approximate P value? | Approximate |
| P value summary | **** |
| Do the medians vary signif. (P < 0.05)? | Yes |
| Number of groups | 4 |
| Kruskal-Wallis statistic | 22.94 |
| Data summary | |
| Number of treatments (columns) | 4 |
| Number of values (total) | 176 |

**Supplementary Table 25.** Multiple comparisons results for Kruskal-Wallis test for GLUT4 Fluorescent Intensity (Groups II,III) data in Figure 5.

| | |
|---|---|
| Number of families | 1 |
| Number of comparisons per family | 6 |
| Alpha | 0.05 |

| Dunn's multiple comparisons test | Mean rank diff. | Significant ? | Summary | Adjusted P Value | |
|---|---|---|---|---|---|
| 1g vs. sμg | 1.279 | No | ns | >0.9999 | A-B |
| 1g vs. 1g | 17.47 | No | ns | 0.6355 | A-C |
| 1g vs. sμg | -33.64 | Yes | * | 0.0111 | A-D |
| sμg vs. 1g | 16.20 | No | ns | 0.8295 | B-C |
| sμg vs. sμg | -34.92 | Yes | ** | 0.0084 | B-D |
| 1g vs. sμg | -51.12 | Yes | **** | <0.0001 | C-D |

| Test details | Mean rank 1 | Mean rank 2 | Mean rank diff. | n1 | n2 | Z |
|---|---|---|---|---|---|---|
| 1g vs. sμg | 84.87 | 83.59 | 1.279 | 46 | 44 | 0.1190 |
| 1g vs. 1g | 84.87 | 67.40 | 17.47 | 46 | 43 | 1.617 |
| 1g vs. sμg | 84.87 | 118.5 | -33.64 | 46 | 43 | 3.113 |
| sμg vs. 1g | 83.59 | 67.40 | 16.20 | 44 | 43 | 1.482 |
| sμg vs. sμg | 83.59 | 118.5 | -34.92 | 44 | 43 | 3.196 |
| 1g vs. sμg | 67.40 | 118.5 | -51.12 | 43 | 43 | 4.652 |

**Supplementary Table 26.** Normality test result for GLUT4 Fluorescent Intensity (Group I) data in Figure 5.

| D'Agostino & Pearson test | | | | |
|---|---|---|---|---|
| K2 | 7.796 | 2.140 | 3.242 | 11.06 |
| P value | 0.0203 | 0.3430 | 0.1977 | 0.0040 |
| Passed normality test (alpha=0.05)? | No | Yes | Yes | No |
| P value summary | * | ns | ns | ** |

**Supplementary Table 27.** ANOVA results for Kruskal-Wallis test for GLUT4 Fluorescent Intensity (Group I) data in Figure 5.

| Table Analyzed | GLUT4 (Group I) |
|---|---|
| Kruskal-Wallis test | |
| P value | 0.0103 |
| Exact or approximate P value? | Approximate |
| P value summary | * |
| Do the medians vary signif. (P < 0.05)? | Yes |
| Number of groups | 4 |
| Kruskal-Wallis statistic | 11.29 |
| Data summary | |
| Number of treatments (columns) | 4 |
| Number of values (total) | 140 |

**Supplementary Table 28.** Multiple comparisons results for Kruskal-Wallis test for GLUT4 Fluorescent Intensity (Group I) data in Figure 5.

| | |
|---|---|
| Number of families | 1 |
| Number of comparisons per family | 6 |
| Alpha | 0.05 |

| Dunn's multiple comparisons test | Mean rank diff. | Significant? | Summary | Adjusted P Value | |
|---|---|---|---|---|---|
| 1g vs. sμg | -24.35 | No | ns | 0.0760 | A-B |
| 1g vs. 1g | 2.886 | No | ns | >0.9999 | A-C |
| 1g vs. sμg | -17.77 | No | ns | 0.4008 | A-D |
| sμg vs. 1g | 27.24 | Yes | * | 0.0299 | B-C |
| sμg vs. sμg | 6.578 | No | ns | >0.9999 | B-D |
| 1g vs. sμg | -20.66 | No | ns | 0.1914 | C-D |

| Test details | Mean rank 1 | Mean rank 2 | Mean rank diff. | n1 | n2 | Z |
|---|---|---|---|---|---|---|
| 1g vs. sμg | 60.89 | 85.24 | -24.35 | 35 | 34 | 2.493 |
| 1g vs. 1g | 60.89 | 58.00 | 2.886 | 35 | 36 | 0.2997 |
| 1g vs. sμg | 60.89 | 78.66 | -17.77 | 35 | 35 | 1.833 |
| sμg vs. 1g | 85.24 | 58.00 | 27.24 | 34 | 36 | 2.808 |
| sμg vs. sμg | 85.24 | 78.66 | 6.578 | 34 | 35 | 0.6736 |
| 1g vs. sμg | 58.00 | 78.66 | -20.66 | 36 | 35 | 2.146 |

**Supplementary Table 29.** Two-way ANOVA test statistical analysis of ARP3 Immunofluorescence data in Figure 6.

| Table Analyzed | ARP3 Immunofluorescence | | | | |
|---|---|---|---|---|---|
| Two-way ANOVA | Ordinary | | | | |
| Alpha | 0.05 | | | | |
| Source of Variation | % of total variation | P value | P value summary | Significant? | |
| Interaction | 0.04768 | 0.9806 | ns | No | |
| Row Factor | 63.43 | <0.0001 | **** | Yes | |
| Column Factor | 0.0001339 | 0.9917 | ns | No | |
| ANOVA table | SS | DF | MS | F (DFn, DFd) | P value |
| Interaction | 112275387 | 2 | 56137694 | F (2, 30) = 0.01958 | P=0.9806 |
| Row Factor | 149353548155 | 2 | 74676774077 | F (2, 30) = 26.05 | P<0.0001 |
| Column Factor | 315320 | 1 | 315320 | F (1, 30) = 0.0001100 | P=0.9917 |
| Residual | 85992683219 | 30 | 2866422774 | | |
| Difference between column means | | | | | |
| Mean of 1g | 359247 | | | | |
| Mean of sμg | 359434 | | | | |
| Difference between means | -187.2 | | | | |
| SE of difference | 17846 | | | | |
| 95% CI of difference | -36634 to 36260 | | | | |
| Data summary | | | | | |
| Number of columns (Column Factor) | 2 | | | | |
| Number of rows (Row Factor) | 3 | | | | |
| Number of values | 36 | | | | |

**Supplementary Table 30.** Multiple comparisons results for Two-way ANOVA test statistical analysis of ARP3 Immunofluorescence data in Figure 6.

| Compare cell means regardless of rows and columns | |
|---|---|
| Number of families | 1 |
| Number of comparisons per family | 15 |
| Alpha | 0.05 |

| Tukey's multiple comparisons test | Mean Diff. | 95.00% CI of diff. | Below threshold? | Summary | Adjusted P Value |
|---|---|---|---|---|---|
| Group I:1g vs. Group I:sμg | 234.9 | -93783 to 94253 | No | ns | >0.9999 |
| Group I:1g vs. Group II:1g | -79527 | -173545 to 14491 | No | ns | 0.1356 |
| Group I:1g vs. Group II:sμg | -75614 | -169632 to 18403 | No | ns | 0.1729 |
| Group I:1g vs. Group III:1g | -155295 | -249313 to -61277 | Yes | *** | 0.0003 |
| Group I:1g vs. Group III:sμg | -160004 | -254022 to -65986 | Yes | *** | 0.0002 |
| Group I:sμg vs. Group II:1g | -79761 | -173779 to 14256 | No | ns | 0.1335 |
| Group I:sμg vs. Group II:sμg | -75849 | -169867 to 18169 | No | ns | 0.1705 |
| Group I:sμg vs. Group III:1g | -155530 | -249548 to -61512 | Yes | *** | 0.0003 |
| Group I:sμg vs. Group III:sμg | -160238 | -254256 to -66221 | Yes | *** | 0.0002 |
| Group II:1g vs. Group II:sμg | 3912 | -90106 to 97930 | No | ns | >0.9999 |
| Group II:1g vs. Group III:1g | -75768 | -169786 to 18249 | No | ns | 0.1713 |
| Group II:1g vs. Group III:sμg | -80477 | -174495 to 13541 | No | ns | 0.1275 |
| Group II:sμg vs. Group III:1g | -79681 | -173699 to 14337 | No | ns | 0.1342 |
| Group II:sμg vs. Group III:sμg | -84389 | -178407 to 9629 | No | ns | 0.0986 |
| Group III:1g vs. Group III:sμg | -4709 | -98726 to 89309 | No | ns | >0.9999 |

| Test details | Mean 1 | Mean 2 | Mean Diff. | SE of diff. | N1 | N2 | q | DF |
|---|---|---|---|---|---|---|---|---|
| Group I:1g vs. Group I:sμg | 280973 | 280739 | 234.9 | 30911 | 6 | 6 | 0.01075 | 30.00 |
| Group I:1g vs. Group II:1g | 280973 | 360500 | -79527 | 30911 | 6 | 6 | 3.638 | 30.00 |
| Group I:1g vs. Group II:sμg | 280973 | 356588 | -75614 | 30911 | 6 | 6 | 3.459 | 30.00 |
| Group I:1g vs. Group III:1g | 280973 | 436268 | -155295 | 30911 | 6 | 6 | 7.105 | 30.00 |
| Group I:1g vs. Group III:sμg | 280973 | 440977 | -160004 | 30911 | 6 | 6 | 7.320 | 30.00 |
| Group I:sμg vs. Group II:1g | 280739 | 360500 | -79761 | 30911 | 6 | 6 | 3.649 | 30.00 |
| Group I:sμg vs. Group II:sμg | 280739 | 356588 | -75849 | 30911 | 6 | 6 | 3.470 | 30.00 |
| Group I:sμg vs. Group III:1g | 280739 | 436268 | -155530 | 30911 | 6 | 6 | 7.116 | 30.00 |
| Group I:sμg vs. Group III:sμg | 280739 | 440977 | -160238 | 30911 | 6 | 6 | 7.331 | 30.00 |
| Group II:1g vs. Group II:sμg | 360500 | 356588 | 3912 | 30911 | 6 | 6 | 0.1790 | 30.00 |
| Group II:1g vs. Group III:1g | 360500 | 436268 | -75768 | 30911 | 6 | 6 | 3.467 | 30.00 |
| Group II:1g vs. Group III:sμg | 360500 | 440977 | -80477 | 30911 | 6 | 6 | 3.682 | 30.00 |
| Group II:sμg vs. Group III:1g | 356588 | 436268 | -79681 | 30911 | 6 | 6 | 3.646 | 30.00 |
| Group II:sμg vs. Group III:sμg | 356588 | 440977 | -84389 | 30911 | 6 | 6 | 3.861 | 30.00 |
| Group III:1g vs. Group III:sμg | 436268 | 440977 | -4709 | 30911 | 6 | 6 | 0.2154 | 30.00 |

**Supplementary Table 31.** Two-way ANOVA test statistical analysis of Glucose Uptake data in Figure 6.

| Table Analyzed | Glucose Uptake | | | |
|---|---|---|---|---|
| Two-way ANOVA | Ordinary | | | |
| Alpha | 0.05 | | | |
| Source of Variation | % of total variation | P value | P value summary | Significant? |
| Interaction | 1.126 | 0.2496 | ns | No |
| Row Factor | 6.916 | 0.0032 | ** | Yes |
| Column Factor | 87.63 | <0.0001 | **** | Yes |
| ANOVA table | SS | DF | MS | F (DFn, DFd) |
| Interaction | 2416747059 | 2 | 1208373529 | F (2, 12) = 1.562 |
| Row Factor | 14840510701 | 2 | 7420255350 | F (2, 12) = 9.590 |
| Column Factor | 188035961946 | 1 | 188035961946 | F (1, 12) = 243.0 |
| Residual | 9284866779 | 12 | 773738898 | |
| Difference between column means | | | | |
| Mean of sµg (control) | 511918 | | | |
| Mean of sµg (+CK666) | 307502 | | | |
| Difference between means | 204416 | | | |
| SE of difference | 13113 | | | |
| 95% CI of difference | 175846 to 232986 | | | |
| Data summary | | | | |
| Number of columns (Column Factor) | 2 | | | |
| Number of rows (Row Factor) | 3 | | | |
| Number of values | 18 | | | |

**Supplementary Table 32.** Multiple comparisons results for Two-way ANOVA test statistical analysis of Glucose Uptake data in Figure 6.

Compare cell means regardless of rows and columns

| | |
|---|---|
| Number of families | 1 |
| Number of comparisons per family | 15 |
| Alpha | 0.05 |

| Tukey's multiple comparisons test | Mean Diff. | 95.00% CI of diff. | Below threshold? | Summary | Adjusted P Value |
|---|---|---|---|---|---|
| Group I:sμg (control) vs. Group I:sμg (+CK666) | 175682 | 99395 to 251969 | Yes | **** | <0.0001 |
| Group I:sμg (control) vs. Group II:sμg (control) | -89179 | -165466 to -12892 | Yes | * | 0.0192 |
| Group I:sμg (control) vs. Group II:sμg (+CK666) | 143255 | 66968 to 219542 | Yes | *** | 0.0004 |
| Group I:sμg (control) vs. Group III:sμg (control) | -75742 | -152029 to 545.2 | No | ns | 0.0520 |
| Group I:sμg (control) vs. Group III:sμg (+CK666) | 129388 | 53101 to 205676 | Yes | ** | 0.0011 |
| Group I:sμg (+CK666) vs. Group II:sμg (control) | -264861 | -341149 to -188574 | Yes | **** | <0.0001 |
| Group I:sμg (+CK666) vs. Group II:sμg (+CK666) | -32427 | -108714 to 43860 | No | ns | 0.7114 |
| Group I:sμg (+CK666) vs. Group III:sμg (control) | -251424 | -327711 to -175137 | Yes | **** | <0.0001 |
| Group I:sμg (+CK666) vs. Group III:sμg (+CK666) | -46294 | -122581 to 29993 | No | ns | 0.3769 |
| Group II:sμg (control) vs. Group II:sμg (+CK666) | 232434 | 156147 to 308722 | Yes | **** | <0.0001 |
| Group II:sμg (control) vs. Group III:sμg (control) | 13437 | -62850 to 89724 | No | ns | 0.9897 |
| Group II:sμg (control) vs. Group III:sμg (+CK666) | 218568 | 142280 to 294855 | Yes | **** | <0.0001 |
| Group II:sμg (+CK666) vs. Group III:sμg (control) | -218997 | -295284 to -142710 | Yes | **** | <0.0001 |
| Group II:sμg (+CK666) vs. Group III:sμg (+CK666) | -13867 | -90154 to 62420 | No | ns | 0.9881 |
| Group III:sμg (control) vs. Group III:sμg (+CK666) | 205130 | 128843 to 281418 | Yes | **** | <0.0001 |

| Test details | Mean 1 | Mean 2 | Mean Diff. | SE of diff. | N1 | N2 | q | DF |
|---|---|---|---|---|---|---|---|---|
| Group I:sμg (control) vs. Group I:sμg (+CK666) | 456944 | 281262 | 175682 | 22712 | 3 | 3 | 10.94 | 12.00 |
| Group I:sμg (control) vs. Group II:sμg (control) | 456944 | 546123 | -89179 | 22712 | 3 | 3 | 5.553 | 12.00 |
| Group I:sμg (control) vs. Group II:sμg (+CK666) | 456944 | 313689 | 143255 | 22712 | 3 | 3 | 8.920 | 12.00 |
| Group I:sμg (control) vs. Group III:sμg (control) | 456944 | 532686 | -75742 | 22712 | 3 | 3 | 4.716 | 12.00 |
| Group I:sμg (control) vs. Group III:sμg (+CK666) | 456944 | 327556 | 129388 | 22712 | 3 | 3 | 8.057 | 12.00 |
| Group I:sμg (+CK666) vs. Group II:sμg (control) | 281262 | 546123 | -264861 | 22712 | 3 | 3 | 16.49 | 12.00 |
| Group I:sμg (+CK666) vs. Group II:sμg (+CK666) | 281262 | 313689 | -32427 | 22712 | 3 | 3 | 2.019 | 12.00 |
| Group I:sμg (+CK666) vs. Group III:sμg (control) | 281262 | 532686 | -251424 | 22712 | 3 | 3 | 15.66 | 12.00 |
| Group I:sμg (+CK666) vs. Group III:sμg (+CK666) | 281262 | 327556 | -46294 | 22712 | 3 | 3 | 2.883 | 12.00 |
| Group II:sμg (control) vs. Group II:sμg (+CK666) | 546123 | 313689 | 232434 | 22712 | 3 | 3 | 14.47 | 12.00 |
| Group II:sμg (control) vs. Group III:sμg (control) | 546123 | 532686 | 13437 | 22712 | 3 | 3 | 0.8367 | 12.00 |
| Group II:sμg (control) vs. Group III:sμg (+CK666) | 546123 | 327556 | 218568 | 22712 | 3 | 3 | 13.61 | 12.00 |
| Group II:sμg (+CK666) vs. Group III:sμg (control) | 313689 | 532686 | -218997 | 22712 | 3 | 3 | 13.64 | 12.00 |
| Group II:sμg (+CK666) vs. Group III:sμg (+CK666) | 313689 | 327556 | -13867 | 22712 | 3 | 3 | 0.8635 | 12.00 |
| Group III:sμg (control) vs. Group III:sμg (+CK666) | 532686 | 327556 | 205130 | 22712 | 3 | 3 | 12.77 | 12.00 |

**Supplementary Table 33.** Unpaired t-test statistical analysis of pAkt/Akt protein expression data in Figure 6.

| Table Analyzed | pAkt/Akt (+ins) |
|---|---|
| Column B | sμg |
| vs. | vs. |
| Column A | 1g |
| Unpaired t test | |
| P value | 0.3887 |
| P value summary | ns |
| Significantly different (P < 0.05)? | No |
| One- or two-tailed P value? | Two-tailed |
| t, df | t=0.9662, df=4 |
| How big is the difference? | |
| Mean of column A | 1.047 |
| Mean of column B | 0.9235 |
| Difference between means (B - A) ± SEM | -0.1239 ± 0.1283 |
| 95% confidence interval | -0.4800 to 0.2322 |
| R squared (eta squared) | 0.1892 |
| F test to compare variances | |
| F, DFn, Dfd | 10.40, 2, 2 |
| P value | 0.1755 |
| P value summary | ns |
| Significantly different (P < 0.05)? | No |
| Data analyzed | |
| Sample size, column A | 3 |
| Sample size, column B | 3 |

**Supplementary Table 34.** Normality test result for GLUT4 Fluorescent Intensity data in Figure 6.

| D'Agostino & Pearson test | | |
|---|---|---|
| K2 | 20.10 | 4.225 |
| P value | <0.0001 | 0.1209 |
| Passed normality test (alpha=0.05)? | No | Yes |
| P value summary | **** | ns |

**Supplementary Table 35.** Mann-Whitney test statistical analysis of GLUT4 Fluorescent Intensity data in Figure 6.

| Table Analyzed | GLUT4 Fluorescent Intensity |
|---|---|
| Column B | +CK-666 |
| vs. | vs. |
| Column A | -CK-666 |
| Mann Whitney test | |
| P value | 0.0004 |
| Exact or approximate P value? | Exact |
| P value summary | *** |
| Significantly different (P < 0.05)? | Yes |
| One- or two-tailed P value? | Two-tailed |
| Sum of ranks in column A,B | 2248 , 1847 |
| Mann-Whitney U | 572 |
| Difference between medians | |
| Median of column A | 3.455, n=40 |
| Median of column B | 2.487, n=50 |
| Difference: Actual | -0.9676 |
| Difference: Hodges-Lehmann | -1.172 |

**Supplementary Table 36.** Normality test result for F-Actin Fluorescent Intensity data in Figure 6.

| D'Agostino & Pearson test | | |
|---|---|---|
| K2 | 10.89 | 16.31 |
| P value | 0.0043 | 0.0003 |
| Passed normality test (alpha=0.05)? | No | No |
| P value summary | ** | *** |

**Supplementary Table 37.** Mann-Whitney test statistical analysis of F-Actin Fluorescent Intensity data in Figure 6.

| Table Analyzed | F-Actin Fluorescent Intensity |
|---|---|
| Column B | +CK-666 |
| vs. | vs. |
| Column A | -CK-666 |
| Mann Whitney test | |
| P value | 0.0308 |
| Exact or approximate P value? | Exact |
| P value summary | * |
| Significantly different (P < 0.05)? | Yes |
| One- or two-tailed P value? | Two-tailed |
| Sum of ranks in column A,B | 2838 , 2212 |
| Mann-Whitney U | 937 |
| Difference between medians | |
| Median of column A | 2.467, n=50 |
| Median of column B | 1.976, n=50 |
| Difference: Actual | -0.4907 |
| Difference: Hodges-Lehmann | -0.5206 |
